# FREQUENT GENE CONVERSION IN HUMAN EMBRYOS INDUCED BY DOUBLE STRAND BREAKS

**DOI:** 10.1101/2020.06.19.162214

**Authors:** Dan Liang, Nuria Marti Gutierrez, Tailai Chen, Yeonmi Lee, Sang-Wook Park, Hong Ma, Amy Koski, Riffat Ahmed, Hayley Darby, Ying Li, Crystal Van Dyken, Aleksei Mikhalchenko, Thanasup Gonmanee, Tomonari Hayama, Han Zhao, Keliang Wu, Jingye Zhang, Zhenzhen Hou, Jumi Park, Chong-Jai Kim, Jianhui Gong, Yilin Yuan, Ying Gu, Yue Shen, Susan B. Olson, Hui Yang, David Battaglia, Thomas O’Leary, Sacha A. Krieg, David M. Lee, Diana H. Wu, P. Barton Duell, Sanjiv Kaul, Jin-Soo Kim, Stephen B. Heitner, Eunju Kang, Zi-Jiang Chen, Paula Amato, Shoukhrat Mitalipov

**Author notes:** These authors contributed equally to this work (Dan Liang, Nuria Marti Gutierrez, Tailai Chen, Yeonmi Lee).

## Abstract

Applications of genome editing ultimately depend on DNA repair triggered by targeted double-strand breaks (DSBs). However, repair mechanisms in human cells remain poorly understood and vary across different cell types. Here we report that DSBs selectively induced on a mutant allele in heterozygous human embryos are repaired by gene conversion using an intact wildtype homolog as a template in up to 40% of targeted embryos. We also show that targeting of homozygous loci facilitates an interplay of non-homologous end joining (NHEJ) and gene conversion and results in embryos which carry identical indel mutations on both loci. Additionally, conversion tracks may expand bidirectionally well beyond the target region leading to an extensive loss of heterozygosity (LOH). Our study demonstrates that gene conversion and NHEJ are two major DNA DSB repair mechanisms in preimplantation human embryos. While gene conversion could be applicable for gene correction, extensive LOH presents a serious safety concern.

## INTRODUCTION

It is believed that DSBs induced by gene editing are typically repaired by two major mechanisms: NHEJ and HDR. Repair by the error-prone NHEJ is dominant and leads to small insertions or deletions (indels), resulting in mutagenic alterations at the cleavage site. In contrast, HDR utilizes an exogenous homologous sequence as a template to repair damaged DNA. NHEJ is active throughout the cell cycle, while HDR is confined to late S/G2 phase (1). While critical for gene therapy applications, frequency of HDR is low with estimated rates of 10-1000-fold lower than those of NHEJ (2–5).

We reported that human preimplantation embryos also employ an alternative HDR mechanism known as gene conversion (6–8). DSBs, selectively induced on a mutant allele in heterozygous embryos, were shown to be frequently repaired by interallelic gene conversion, utilizing the homologous wildtype (WT) allele as a template. Remaining DSBs tended to be resolved by NHEJ, with HDR rarely occurring.

Gene conversion is a process of a unidirectional copy of the genetic code from a highly homologous donor DNA template to an acceptor sequence, and is typically initiated as a repair response to DSB on the acceptor DNA (9). Repair by gene conversion is activated from an intact template DNA, such as parental allele on homologous chromosome (interallelic) or other highly homologous sequences on the same chromosome (interlocus). It is suggested that gene conversion exploits the synthesis-dependent strand annealing (SDSA) that includes 5’→3’ resection, followed by homologous strand invasion and DNA synthesis (10, 11). As a result, the DSB locus and adjacent area become identical to the template DNA, leading to one of the hallmarks of gene conversion, that is acquisition of homozygosity at the target region, or LOH. Gene conversion typically occurs as meiotic non-crossover recombination, while the frequency of mitotic gene conversion is much lower (300-1000 fold) (12, 13). The extent of the sequence region that is copied from the donor to acceptor, known as conversion tract length, in meiotic cells is generally short, reaching a few hundred base pairs (bp). In contrast, mitotic conversion tracts often extend substantially, both upstream and downstream, from DSB loci, and leads to more extensive LOH (14).

Repair outcomes of NHEJ and HDR with an exogenous template can be easily identified by detection of novel indel mutations or marker SNPs in bulk DNA of pooled cells from whole mammalian embryos. In contrast, gain of homozygosity caused by gene conversion is more difficult to recognize, particularly if homozygous loci are targeted. In addition, due to mixture of all parental alleles with various DSB repair outcomes and mosaicism in mammalian embryos, gene conversion can be easily overlooked in pooled DNA samples unless single-cell analyses are performed.

By inducing DSBs at various heterozygous and homozygous loci and accurately analyzing LOH at the single-cell level, we show here that gene conversion is a common DNA repair pathway in human embryos with frequencies comparable to those of NHEJ. In homozygous loci, DSBs repair by gene conversion is often combined with NHEJ or HDR within the same cell, leading to copying of repair outcomes from one chromosome to another.

## RESULTS

### Gene conversion frequency at heterozygous loci

In an effort to determine the frequency of DNA repair by interallelic gene conversion, we generated heterozygous human zygotes by fertilization of WT metaphase II (MII) oocytes with sperm donated by a subject carrying a heterozygous mutation (1 bp C>T substitution; g.15819 C>T, NG_007884.1) in exon 22 of *MYH7* gene located on chromosome 14, and implicated in familial hypertrophic cardiomyopathy (HCM). We introduced the sgRNA targeting the mutant paternal *MYH7* allele along with Cas9 protein and exogenous single-stranded oligodeoxynucleotide (ssODN) into cytoplasm of pronuclear stage zygotes 18 hours after fertilization (**Fig. S1A**). Injected zygotes (N=86) along with non-injected controls (N=18) were cultured for 3 days and then cleaving 4-8 cell stage embryos were disaggregated, and each blastomere was individually analyzed by Sanger sequencing.

As expected for heterozygous (*MYH7^WT/Mut^*) sperm, on-target analysis of individual blastomeres (N=110) disaggregated from 18 control embryos revealed that 9 were uniformly heterozygous *MYH7^WT/Mut^* with every blastomere showing the same genotype. Individual blastomeres from the remaining controls presented only WT sequences, indicating that these embryos were uniformly homozygous (*MYH7^WT/WT^*) as a result of fertilization with WT sperm **(Fig. 1A)**. In contrast to controls, no uniform*MYH7^WT/Mut^* heterozygous embryos with an intact g.15819 C>T mutation were discovered among injected embryos. Every blastomere (N=515) analyzed from 86 injected embryos contained an intact WT *MYH7* allele suggesting absence of mistargeting of the WT allele and exceptional fidelity of the selected sgRNA. Majority of injected embryos (58/86, 67.4%) were uniformly homozygous with each sister blastomere showing WT *MYH7* allele only **(Fig. 1A).** While it is possible that these homozygous WT embryos originated from the WT sperm, increase in the portion of *MYH7^WT/WT^* embryos compared to controls was similar to our previous observations with *MYBPC3* mutation (7). Among the remaining embryos, 6/86 (7.0%) were uniformly heterozygous carrying the WT and indel mutation at or adjacent to the pre-existing mutant locus (*MYH7^WT/Indel^).* Note, that all sister blastomeres in these embryos carried identical indel mutations. The rest of embryos (22/86; 25.6%) were mosaic, each consisting of blastomeres with mixed *MYH7^WT/Mut^, MYH7^WT/Indel^* and *MYH7^WT/WT^* genotypes **(Fig. 1A)**. Remarkably, no HDR with ssODN was found in blastomeres (N=515) of injected embryos.

**Fig 1.**
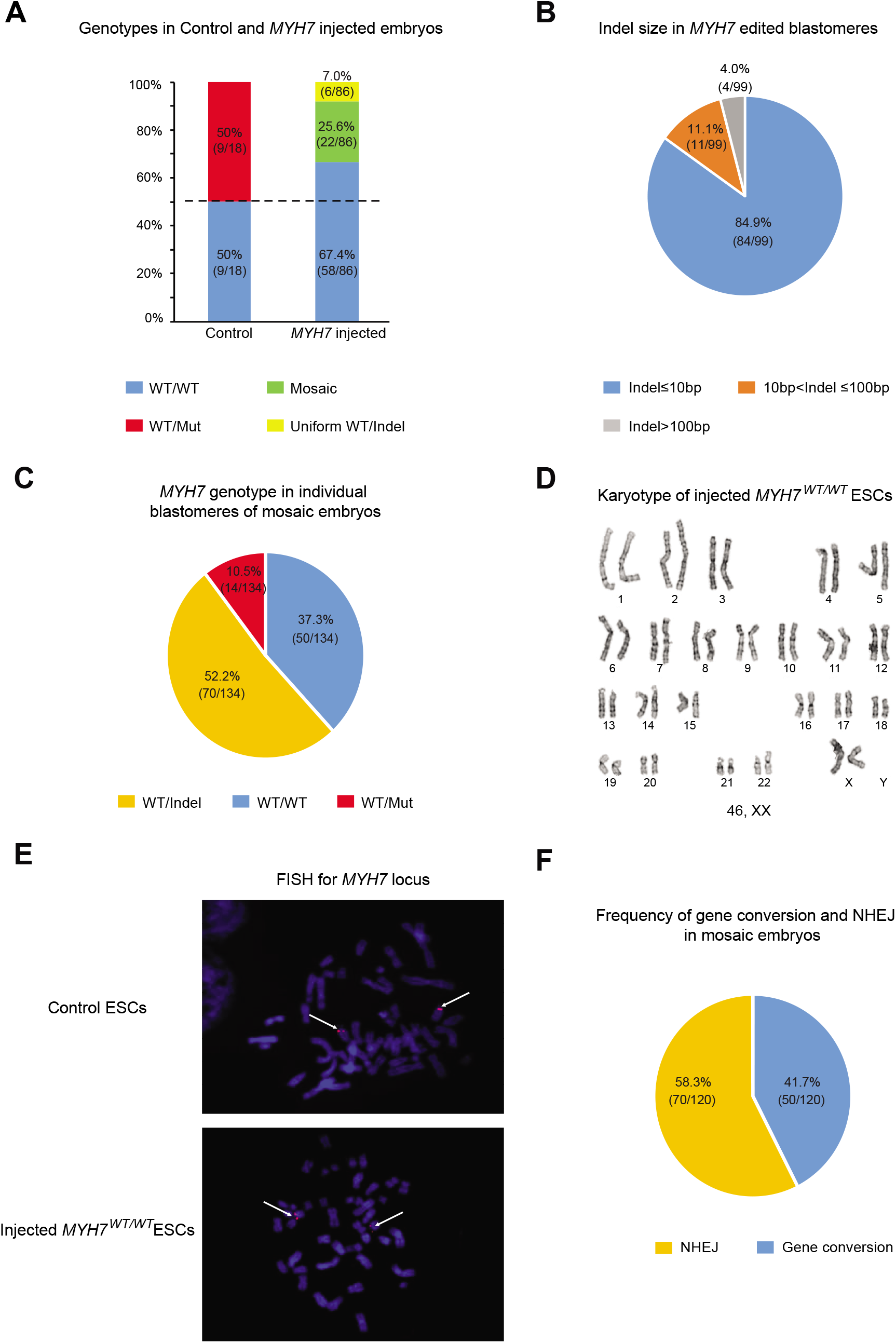
DNA DSB repair outcomes in human embryos heterozygous for *MYH7* locus. (**A**) *MYH7* target region genotype outcomes in control and injected human embryos. Half of the control embryos were *MYH7^WT/WT^* homozygous while other half were *MYH7^WT/Mut^* heterozygous, typical for sperm from the heterozygous subject. (**B**) Indel size distribution in individual blastomeres with *MYH7^WT/Indel^* genotypes from uniform WT/indel and mosaic embryos demonstrates that the majority of mutations (85%) are small indels (≤10 bp). (**C**) *MYH7* target region genotypes outcomes in individual blastomeres of mosaic embryos. (**D**) G-banding analysis of *MYH7^WT/WT^* ESCs derived from injected blastocysts exhibited normal euploid karyotype. (**E**) FISH labeling of the *MYH7* locus in *MYH7^WT/WT^* ESC lines. Strong signals (arrows) are seen on both parental chromosome 14 indicating the presence of two intact *MYH7* alleles. (**F**) Frequency of gene conversion and NHEJ in edited blastomeres of mosaic embryos.

Since at least one blastomere in every mosaic embryo carried either intact g.15819 C>T mutant locus or indel mutation, we presumed that these embryos were fertilized with the mutant sperm. Among 99 *MYH7^WT/Indel^* blastomeres from uniform WT/Indel and mosaic embryos (N=28), majority of indels (84.9%, 84/99) were represented as small insertions, deletions or substitutions (≤10 bp), while 11.1% (11/99) were medium size (11-100 bp). Only 4 (4.0%) blastomeres (all from one mosaic embryo) contained a large 652 bp deletion (**Fig. 1B)**.

In-depth analysis of 134 blastomeres isolated from 22 mosaic embryos revealed that 14 (10.5%) were heterozygous with a WT and an intact mutant allele (*MYH7^WT/Mut^*) while 70 (52.2%) were heterozygous with WT and indel mutations (*MYH7^WT/Indel^*) (**Fig. 1C**). Consistent with our previous studies, the remaining blastomeres (50/134, 37.3%) lost the mutant allele and appeared as homozygous *MYH7^WT/WT^*. To eliminate the possibility of large deletions on the mutant paternal *MYH7* allele, we reanalyzed all *MYH7^WT/WT^* blastomeres from mosaic and uniform *MYH7^WT/WT^* embryos with several long-range PCR primers amplifying from 2301 bp to 8190 bp fragments surrounding the *MYH7* g.15819 C>T mutant locus. No evidence of large deletions was found by detailed analysis of gel bands (**Fig. S1B**).

To exclude potential cases of allele dropout during whole genome amplification (WGA) of single blastomere DNA, we validated our results on ESCs derived from injected embryos. We established a total of 14 ESC lines from injected blastocysts that provided unlimited amount of DNA for more detailed analyses (without WGA). On-target genotyping demonstrated that 8 cell lines were homozygous *MYH7^WT/WT^*, 3 were heterozygous *MYH7^WT/Indel^* and remaining 3 were intact heterozygous *MYH7^WT/Mut^.* Detailed G-banding analysis confirmed that all ESC lines carried normal euploid karyotypes without any detectable deletions or other cytogenetic abnormalities (**Fig. 1D**). Moreover, Short Tandem Repeat (STR) analysis excluded the possibility of parthenogenesis and confirmed paternity of the *MYH7* sperm carrier in all cell lines (**Table S1**). Next, fluorescence in situ hybridization (FISH) assay was employed to detect and visualize presence of *MYH7* alleles within individual nuclei. The FISH probe was designed to hybridize precisely to the *MYH7* target locus. All *MYH7^WT/WT^* cell lines including controls demonstrated the presence of two signals in each nucleus consistent with the conclusion that all *MYH7^WT/WT^* cells indeed carry two intact alleles (**Fig. 1E**).

Taken together, our results suggest that a large percentage of DSBs (41.7%, 50/120), are resolved by gene conversion. Its frequency is comparable to that of repair by NHEJ (58.3%, 70/120) (**Fig. 1F)**. Remarkably, HDR via exogenous ssODN was not employed. These conclusions are consistent with our previous study with human heterozygous *MYBPC3* embryos (7), and indicate that gene conversion is one of the major DNA DSB repair pathways in human heterozygous embryos.

### Conversion tract and LOH in human embryos

Typically, gene conversion induces LOH not only within the target locus but also in adjacent upstream and downstream heterozygous sites (8). Therefore, we screened blood DNA from egg donors and the *MYH7* sperm donor and identified single nucleotide polymorphisms (SNPs) differentiating parental alleles in embryos. A total of 13 informative SNPs was identified between the egg donor 1 and the *MYH7* sperm donor that were distributed at various distances upstream and downstream from the *MYH7* locus on chromosome 14 (**Fig. 2A and Table S2**). Of these, 8 SNP positions were homozygous in egg donor 1, but heterozygous in the sperm donor DNA. Analysis of control *MYH7^WT/Mut^* embryos indicated that the mutant allele is genetically linked with unique proximal nucleotides at these SNPs, suggesting lack of meiotic recombination at those loci.

**Fig 2.**
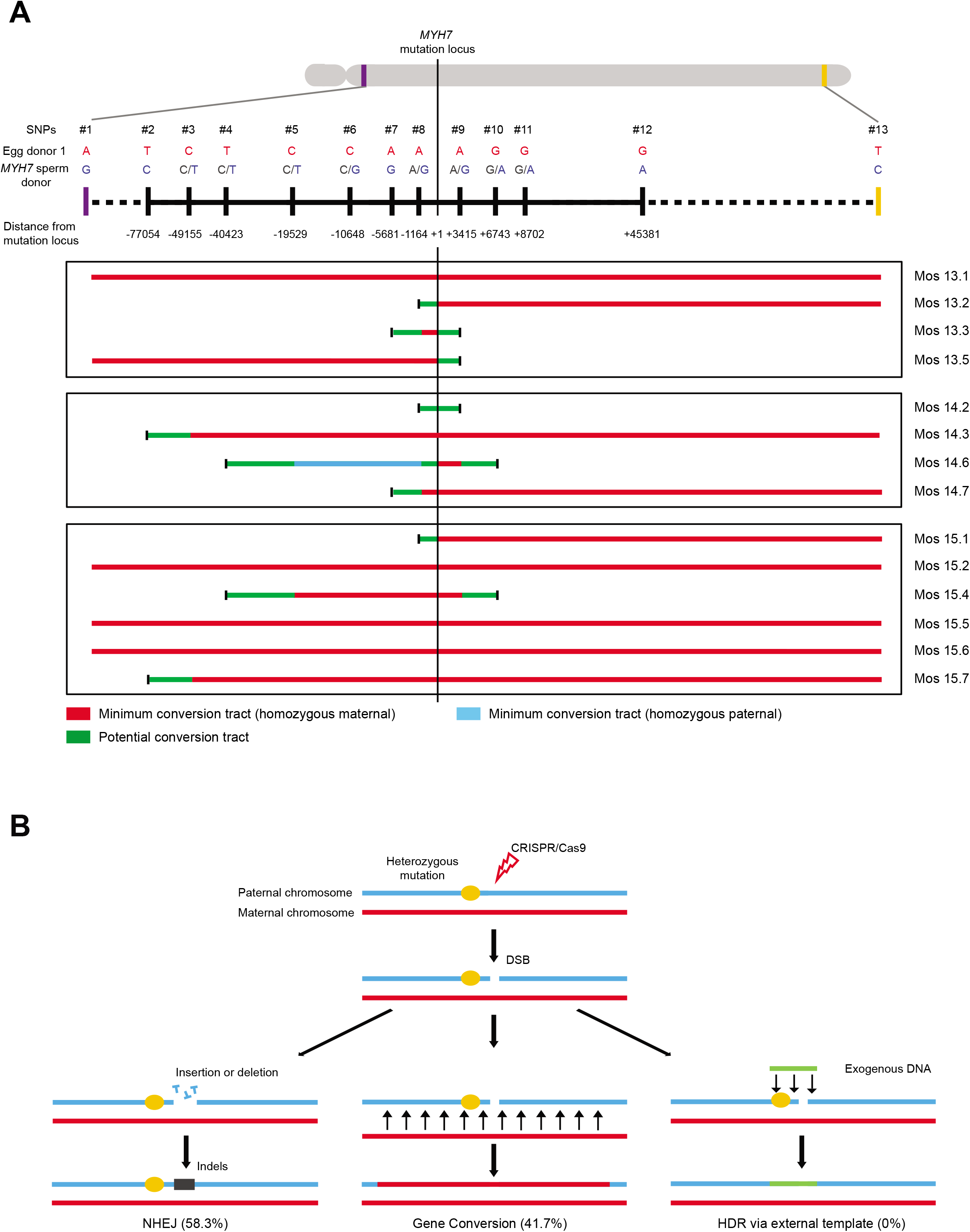
Gene conversion tract and frequency in mosaic *MYH7* embryos. (**A**) LOH induced by long gene conversion tracts. Schematic map of 13 informative SNPs located upstream and downstream at various distances from the mutant *MYH7* locus differentiating parental alleles in individual blastomeres from mosaic embryos. LOH at each SNP site was determined in sister blastomeres with *MYH7^WT/WT^* genotype from 3 mosaic embryos and compared to *MYH7^WT/Indel^* and intact control blastomeres. LOH and direction of conversion tract in each blastomere is indicated in color lines. Red lines indicate that these loci lost paternal nucleotides and become homozygous maternal. Light blue lines indicate that these loci lost maternal contribution and became homozygous paternal. Green lines indicate that these loci are homozygous but LOH cannot be determined because of initial heterozygosity in contributing gametes. Black bars at the end of lines indicate that these loci are heterozygous. See for more details in **Table S2**. (**B**) Schematic summary of DSB repair pathways in human heterozygous embryos. Selective induction of DSBs on the mutant paternal allele is repaired either by NHEJ or gene conversion. HDR via exogenous DNA template is not observed.

We genotyped these loci in all *MYH7^WT/WT^* and *MYH7^WT/Indel^* blastomeres from three mosaic embryos (MYH7-Mos13, 14 and 15 in **Fig. 2A**) produced from the egg donor 1 and the sperm donor, and compared these to controls. As expected, *MYH7^WT/Mut^* blastomeres in controls and *MYH7^WT/Indel^* blastomeres from mosaic embryos were heterozygous at all SNP loci. However, almost all *MYH7^WT/WT^* blastomeres from injected embryos were homozygous at some or most SNP positions, indicating LOH consistent with gene conversion (**Fig. 2A and Table S2**). The direction and length of the conversion tract differed, even among sister blastomeres. For instance, blastomere Mos13.2 retained heterozygous loci upstream of SNP8 (−1164 bp, A/G), while all loci downstream of SNP8 were homozygous carrying exclusively maternal nucleotides. In contrast, its sister blastomere Mos13.5 showed conversion tract in the opposite direction with LOH expanding upstream of SNP9 (+3415 bp, A/G). Blastomere 13.3 showed the shortest conversion tract length of 1164 bp, while LOH in other three sister blastomeres extended beyond examined SNPs. While in most LOH cases we detected maternal nucleotides indicating that the maternal allele was used as a template, one blastomere Mos14.6 showed presence of paternal nucleotides at SNP5, 6, 7 and 8 loci (T/T, G/G, G/G and G/G, respectively) (**Fig. 2A and Table S2**). This could be caused by two DSBs on the paternal genome, and the combined effect of crossover and gene conversion. The first DSB could result in a crossover followed by gene conversion after second DSB.

As already shown, the percentage of uniform *MYH7^WT/WT^* embryos was substantially increased (67.4%) in the injected group compared to expected 50%. Unlike mosaic embryos, the sperm origin (mutant or WT) in uniform *MYH7^WT/WT^* embryos could not be determined, making it difficult to evaluate contribution of gene conversion in this group. However, since gene conversion is associated with substantial LOH, we reanalyzed expected heterozygous loci adjacent to the target site in uniform *MYH7^WT/WT^* embryos derived from the egg donor 1. The SNP7, adjacent to the mutant locus, was A/A homozygous in egg donor 1 but G/G homozygous in the sperm donor (**Fig. S2A**). At the same time, this egg donor was C/C homozygous at SNP6 locus but the sperm donor was heterozygous (C/G). In control embryos, we determined that the mutant paternal allele was always linked with G nucleotide at this locus. Analysis of individual blastomeres from 11 uniform *MYH7^WT/WT^* injected embryos produced from the egg donor 1 revealed that 8 embryos were all uniformly heterozygous (A/G) at SNP7 and homozygous (C/C) at SNP6 loci indicating that these embryos were likely fertilized by the WT sperm (MYHY-WT21.1 in **Fig. S2B and Table 1**). However, in the remaining 3 embryos (MYH7-WT22, 27 and 31 in **Table 1**) some blastomeres become homozygous at SNP7 (A/A), carrying maternal nucleotides while other sister blastomeres from the same embryos were heterozygous at this site. At the SNP6, some blastomeres from these embryos were still heterozygous (C/G) while other sister blastomeres become (C/C) homozygous (**Fig. S2B and Table 1**). Since the G locus is linked to the mutant *MYH7* locus, thse embryos were likely fertilized by the mutant sperm but later repaired by gene conversion

**Table 1.** LOH at adjacent to *MYH7* heterozygous loci in individual blastomeres

| **Embryo ID** | **Blastomere ID** | **MYH7 on-target genotype** | **SNP6 genotype (rs41285540, - 10648 bp)** | **SNP7 genotype (rs3729820, - 5681 bp)** |
| --- | --- | --- | --- | --- |
| Egg donor 1 | n/a | WT/WT | C | A |
| Sperm donor | n/a | WT/Mut | C/G | G |
| Control 12 | 12.1 | WT/Mut | C/G | A/G |
| Control 13 | 13.1 | WT/Mut | C/G | A/G |
| MYH7- WT21 | 21.1 | WT/WT | C | A/G |
|  | 21.2 | WT/WT | C | A/G |
| MYH7- WT23 | 23.1 | WT/WT | C | A/G |
|  | 23.2 | WT/WT | C | A/G |
| MYH7- WT24 | 24.1 | WT/WT | C | A/G |
|  | 24.2 | WT/WT | C | A/G |
| MYH7- WT25 | 25.1 | WT/WT | C | A/G |
|  | 25.2 | WT/WT | C | A/G |
| MYH7- WT26 | 26.1 | WT/WT | C | A/G |
|  | 26.2 | WT/WT | C | A/G |
| MYH7- WT28 | 28.1 | WT/WT | C | A/G |
|  | 28.2 | WT/WT | C | A/G |
| MYH7- WT29 | 29.1 | WT/WT | C | A/G |
|  | 29.2 | WT/WT | C | A/G |
| MYH7- WT30 | 30.1 | WT/WT | C | A/G |
|  | 30.2 | WT/WT | C | A/G |
| MYH7- WT22 | 22.1 | WT/WT | C | A |
|  | 22.2 | WT/WT | C | A |
|  | 22.3 | WT/WT | C | A |
|  | 22.4 | WT/WT | C/G | A/G |
|  | 22.5 | WT/WT | C/G | A/G |
|  | 22.6 | WT/WT | C | A |
|  | 22.7 | WT/WT | C/G | A/G |
|  | 22.8 | WT/WT | C | A |
| MYH7- WT27 | 27.1 | WT/WT | C | A |
|  | 27.2 | WT/WT | C | A |
|  | 27.3 | WT/WT | C | A |
| MYH7- WT31 | 31.1 | WT/WT | C | A |
|  | 31.2 | WT/WT | C | A |
|  | 31.3 | WT/WT | C | A |
|  | 31.4 | WT/WT | C/G | A/G |
|  | 31.5 | WT/WT | C | A |
|  | 31.6 | WT/WT | C/G | A/G |
|  | 31.7 | WT/WT | C/G | A/G |
|  | 31.8 | WT/WT | C | A |

Based on these results, it is reasonable to assume that these 3 *MYH7^WT/WT^* embryos were fertilized by the mutant sperm but were subsequently repaired by gene conversion. In summary, DSB repair by gene conversion in human embryos is associated with bidirectional or unidirectional conversion tracts ranging from minimum of 1164 bp to maximum expanding the entire chromosome arm and leading to a substantial LOH. High frequency of gene conversion leads to increased yield of uniform *MYH7^WT/WT^* embryos derived by fertilization with heterozygous sperm. Based on these results we estimate that actual gene conversion rates could be much higher than that calculated from mosaic embryos alone (**Fig. 2B)**.

### High frequency of homozygosity induced by DSBs

To evaluate if HDR via ssODN can be achieved when inducing DSBs at homozygous loci we targeted the WT homozygous *MYBPC3* locus (g.14846, NG_007667.1) in human embryos (**Fig. S3A**). CRISPR/Cas9 along with ssODN was injected into the cytoplasm of 32 WT oocytes during fertilization with WT sperm (M-phase) and 21 pronuclear stage zygotes 18 hours after fertilization (S-phase). Each blastomere (N=321) in cleaving embryos was individually isolated and the target DNA locus was amplified by long-range PCR primers (1742 bp, 3054 bp and 8424 bp fragment size in **Fig. S3B**).

Sanger sequencing revealed that a large proportion of injected blastomeres (40.2%, 129/321) lost both WT alleles but showed the presence of only one indel mutation (designated as *MYBPC3^homo-Indel/Indel^*). Remarkably, a few blastomeres (5.9%, 19/321) carried a sequence identical to the ssODN on one or both alleles (*MYBPC3^WT/HDR^, MYBPC3^Indel/HDR^* or *MYBPC3^HDR/HDR^*) indicating HDR with the exogenous template. In addition, 26 blastomeres (8.1%) showed the WT allele only and were deemed as non-targeted *MYBPC3^WT/WT^* (**Fig. 3A**). We also found that a small portion of blastomeres (8.7%, 28/321) carried one intact WT allele and one indel mutation (classified as *MYBPC3^WT/Indel^).* Remaining blastomeres (37.1%, 119/321) presented two different indel mutations and were designated as compound heterozygous *MYBPC3^Indel/Indel^* (**Fig. 3A**).

**Fig 3.**
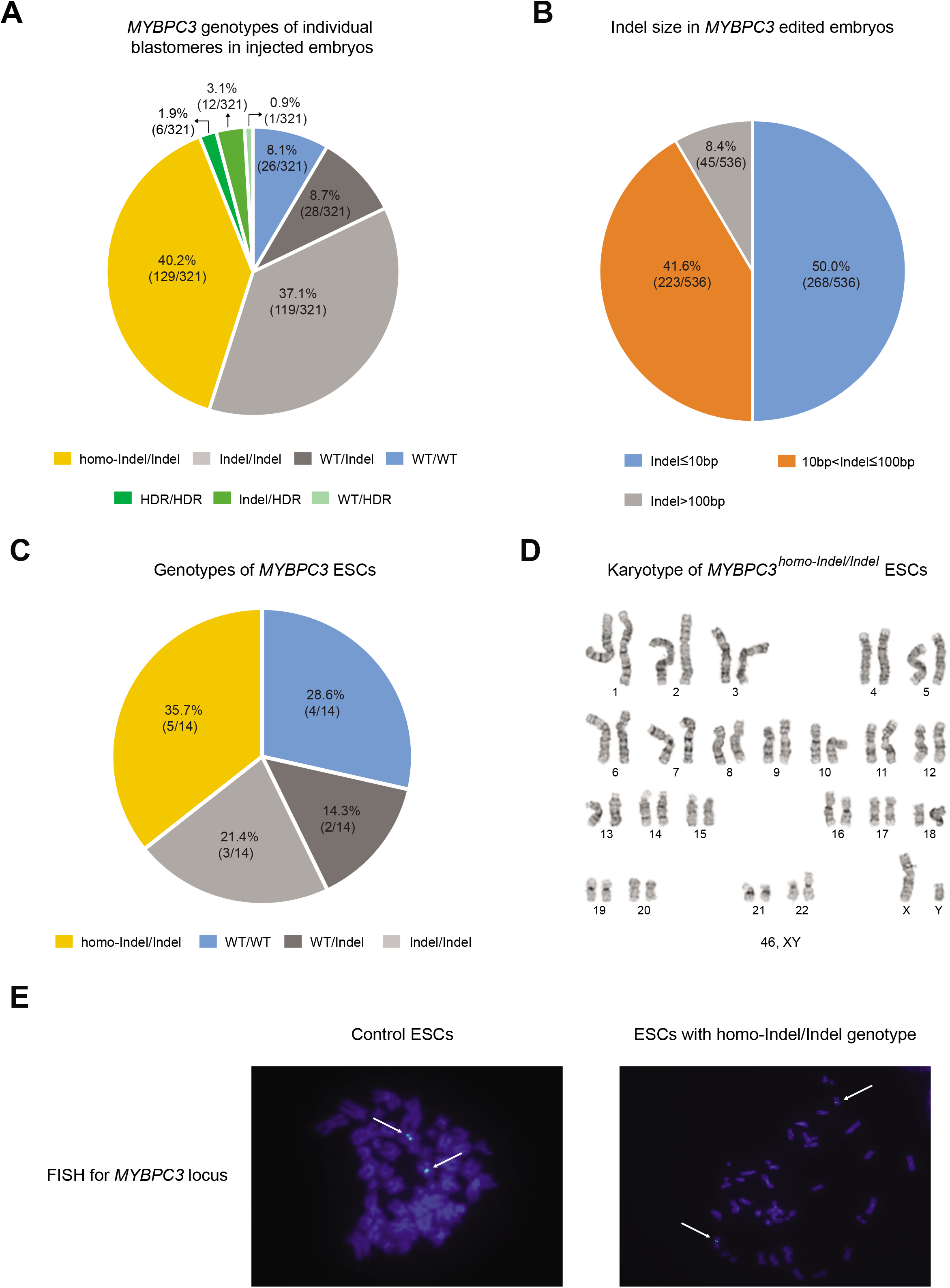
Gene conversion at homozygous *MYBPC3* locus. (**A**) On-target *MYBPC3* genotypes in individual blastomeres of human embryos injected with CRISPR/Cas9. Note that a large proportion of blastomeres carried identical indels (homo-Indel/Indel) indicating gene conversion. In contrast to heterozygous loci, DSB repair in homozygous *loci* may be resolved by HDR but at low frequency. (**B**) Indel size in blastomeres repaired by NHEJ. (**C**)*MYBPC3* genotypes of 14 ESC lines derived from CRISPR/Cas9 injected embryos. Similar to that seen in blastomeres, a substantial number of ESC lines carried identical (homo-Indel/Indel) indels. (**D**) G-banding analysis confirming that all ESC lines carrying identical indels (*MYBPC3^homo-Indel/Indel^*) exhibit normal diploid karyotypes without any detectable large deletions. (**E**) FISH labeling of the *MYBPC3* locus in ESC lines. Strong signals (arrows) are seen on both parental chromosomes indicating the presence of two intact *MYBPC3* alleles in all *MYBPC3^homo-Indel/Indel^* cell lines.

Among all indel mutations (N=536), half (268/536) carried small mutations (≤ l0 bp), 41.6% (223/536) were medium size (10-100 bp) while the remaining 8.4% (45/536) were over >100 bp including several large deletions ranging in the size up to 3.8kb (**Fig. 3B**).

To exclude the possibility of even larger deletions or allele dropouts, we derived 14 ESC lines from treated blastocysts and carried out more detailed analyses. Five cell lines (35.7%) were genotyped as *MYBPC3^homo-Indel/Indel^*, the remaining were either *MYBPC3^Indel/Indel^, MYBPC3^WT/Indel^* or *MYBPC3^WT/WT^* (**Fig. 3C)**. G-banding cytogenetic assay and STR analysis indicated that all 14 cell lines had normal euploid karyotype and were derived by fertilization, not parthenogenesis (**Fig. 3D and Table S3**). Labeling with FISH probes designed to bind to the target *MYBPC3* locus demonstrated two fluorescent signals within each cell, ruling out the possibility of large deletions or allele dropouts (**Fig. 3D**). Thus, it is reasonable to conclude that blastomeres with *MYBPC3^homo-Indel/Indel^* or *MYBPC3^HDR/HDR^* genotypes are indeed homozygous.

### Gene conversion and LOH at homozygous loci

We reasoned that due to the random nature of NHEJ, the chances of generating identical indel mutations on both alleles are very low. However, as we indicated above, 40.2% of blastomeres (**Fig. 3A**) carried identical indels. We postulated that during fertilization and zygotic stages, parental alleles may not be equally accessible for CRISPR/Cas9. Such a scenario would lead to initial targeting one of the parental alleles and generating an indel mutation on the oocyte or sperm allele first. Later, the second allele becomes available and targeted by CRISPR/Cas9, but DSB repair would now be resolved by gene conversion leading to copying of the first indel mutation to the second allele. Therefore, we hypothesized that these results may represent a sequential repair by NHEJ and gene conversion.

To test this assumption, we investigated LOH at adjacent heterozygous loci as a result of gene conversion. We used two informative SNPs (SNP14 and 15) located next to the target site to differentiate egg donors 1 and 2 from the sperm alleles (**Fig. 4A**). Since *both MYBPC3* alleles were identical, we also asked which of the parental alleles was used as a template for gene conversion. As expected, intact control *MYBPC3^WT/WT^* and compound heterozygous *MYBPC3^Indel/Indel^* blastomeres (MYBPC3 S11.6 in **Fig. 4B**, MYBPC3 M31.1 in **Fig. 4D)** derived from egg donor 1 (G/G at SNP14) and egg donor 2 (G/G at both SNP14 and 15) and the sperm donor (A/A at SNP14 and C/C at SNP15) were heterozygous at these loci.

**Fig 4.**
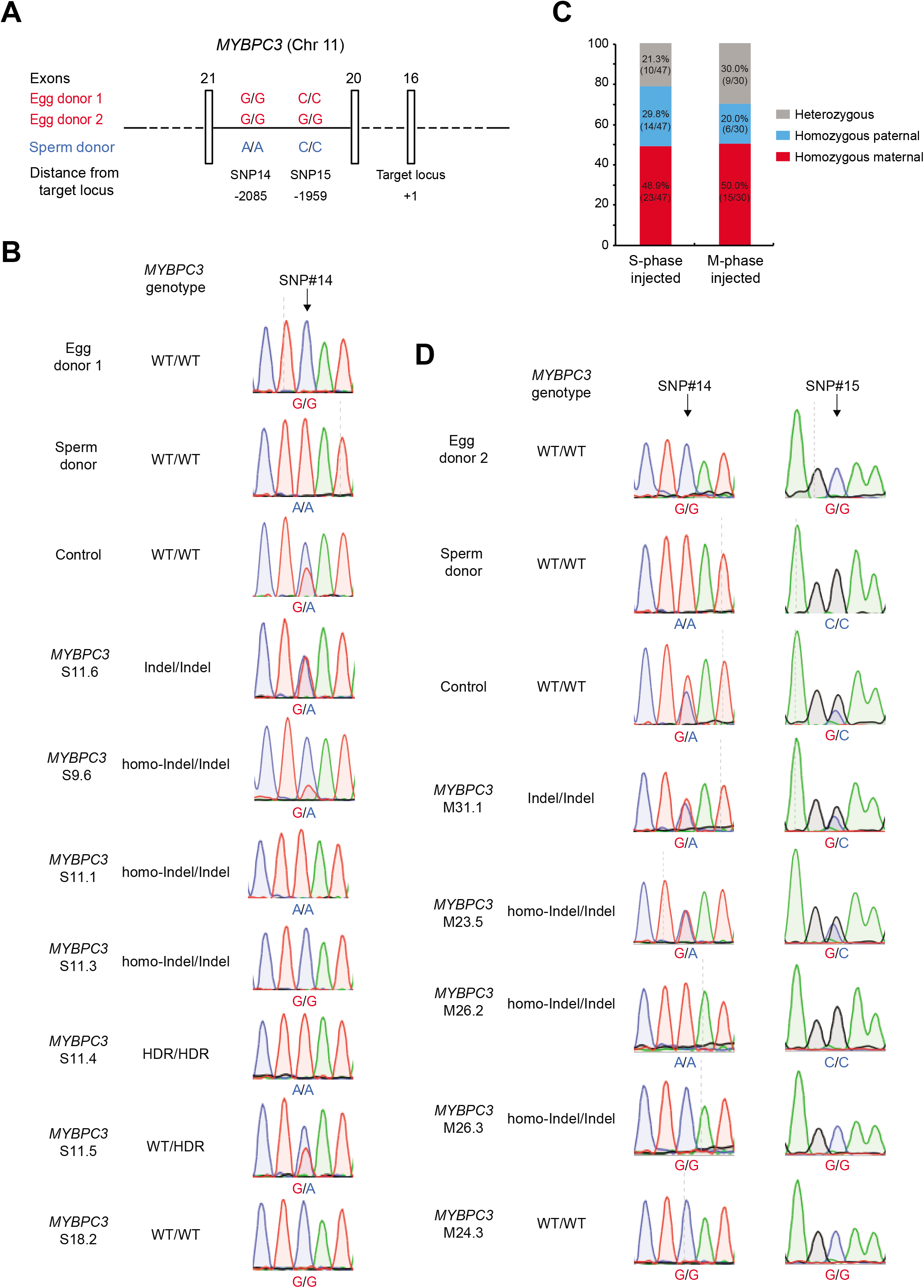
LOH due to gene conversion in *MYBP3* embryos. (**A**) Schematic map of two heterozygous loci (SNP14 and 15) located upstream of target *MYBPC3* locus in gamete donors. Boxes indicate exons. (**B**) Sanger sequencing chromatographs showing SNP14 genotypes in individual blastomeres from control and edited embryos generated from egg donor 1 and *MYBPC3* WT sperm donor. Note that edited blastomeres with *MYBPC3^homo-Indel/Indel^, MYBPC3^HDR/HDR^* and *MYBPC3^WT/WT^* genotypes lost heterozygosity at SNP14 locus. (**C**) LOH due to gene conversion in *MYBPC3^homo-Indel/Indel^* blastomeres from S-phase and M-phase injected embryos. Note that in both groups the maternal allele is preferentially used as a template. (**D**) Sanger sequencing chromatographs showing LOH at SNP14 and 15 loci in individual blastomeres with *MYBPC3^homo-Indel/Indel^* and *MYBPC3^WT/WT^* genotypes from embryos of the egg donor 2 and sperm donor.

However, 78.7% (37/47) blastomeres with *MYBPC3^homo-Indel/Indel^* genotype derived from egg donor 1 lost heterozygosity at the SNP14 locus (S-phase injection) (**Table S4**). Most of these LOH blastomeres (23/37, 62.2%) retained the maternal genotype (G/G, MYBPC3 S11.3 in **Fig. 4B**), while 14 (37.8%) carried paternal (A/A) allele (MYBPC3 S11.1 in **Fig. 4B**). The remaining *MYBPC3^homo-Indel/Indel^* blastomeres (10/47, 21.3%) were heterozygous at this locus (MYBPC3 S9.6 in **Fig. 4, B and C**).

In the M-phase group, 70.0% (21/30) of blastomeres with *MYBPC3^homo-Indel/Indel^* genotype derived from egg donor 2 lost heterozygosity at both SNP14 and 15 loci (**Table S5**). Similar to the S-phase, the majority of these LOH blastomeres (15/21, 71.4%) retained maternal nucleotides (MYBPC3 M26.3 in **Fig. 4D**), while the remaining (6/21, 28.6%) carried paternal nucleotides (MYBPC3 M26.2 in **Fig. 4D**). The rest of *MYBPC3^homo-Indel/Indel^* blastomeres (9/30, 30.0%) were heterozygous at both SNP14 and 15 loci (MYBPC3 M23.5 in **Fig. 4D)**, likely due to shorter conversion tracks (**Fig. 4C**).

Postulating that homozygosity and LOH are the hallmarks of gene conversion, we next genotyped other homozygous groups, i.e. *MYBPC3^WT/WT^* and *MYBPC3^HDR/HDR^*. All 3 *MYBPC3^WT/WT^* blastomeres derived from egg donor 1 lost heterozygosity at the informative SNP14 locus and retained the maternal nucleotides (MYBPC3 S18.2 in **Fig. 4B**). Likewise, two of three *MYBPC3^WT/WT^* blastomeres produced from egg donor 2 also became homozygous at both SNP14 and 15 loci suggestive gene conversion (MYBPC3 M24.3 in **Fig. 4D**). Moreover, 3 of 4 analyzed *MYBPC3^HDR/HDR^* blastomeres derived from egg donor 1 also become homozygous at the informative SNP14 locus but retained paternal genotype (MYBPC3 S11.4 in **Fig. 4B**). These observations suggest that the majority of blastomeres with homozygous genotypes at the targeted locus (*MYBPC3^WT/WT^* or *MYBPC3^HDR/HDR^*) are also the result of gene conversion.

Taken together, these results confirm LOH adjacent to the *MYBPC3* locus and suggest that many homozygous blastomeres are likely generated by interallelic gene conversion. It is facilitated by asynchronous induction of DSBs on parental alleles leading to generation of an indel mutation or HDR. Subsequent DSB on the second allele activates gene conversion resulting in copying of the indel mutation or HDR to both alleles. Moreover, DSBs occur preferentially on the oocyte allele first irrespective of M-phase or S-phase injections. It is possible, that sperm alleles are less accessible for DSBs during early post-fertilization stages of development in human embryos.

### Gene conversion frequency at homozygous loci

To accurately estimate gene conversion frequency, it is critical to validate LOH at heterozygous sites located close to the target locus. However, if the conversion tract is short, distant SNPs are often not informative to account for all cases of LOH. In an effort to further corroborate gene conversion, we recruited a sperm donor homozygous for *LDLRAP1* (g.24059 G>A, NG_008932.1) mutation located on chromosome 1 and associated with familial hypercholesterolemia (FH). The sgRNA was designed to target the wildtype locus immediately upstream of the *LDLRAP1* mutation site (**Fig. 5A**). CRISPR/Cas9 was co-injected with homozygous mutant *LDLRAP1* sperm into the cytoplasm of WT MII oocytes (N=19). Injected zygotes along with intact controls (N=2) were cultured for 3 days and individual blastomeres were analyzed as described above. As expected, all 9 blastomeres derived from 2 control embryos were uniformly heterozygous at the mutant locus (A/G; *LDLRAP1^WT/Mut^*). Among 19 injected embryos, only 4 (21.1%) were uniformly *LDLRAP1^WT/Mut^* heterozygous and were thus regarded as non-targeted (**Fig. 5B**). However, one embryo lost heterozygosity in all blastomeres and became uniformly *LDLRAP1^WT/WT^* indicating repair by gene conversion using the WT maternal allele as a template.

**Fig 5.**
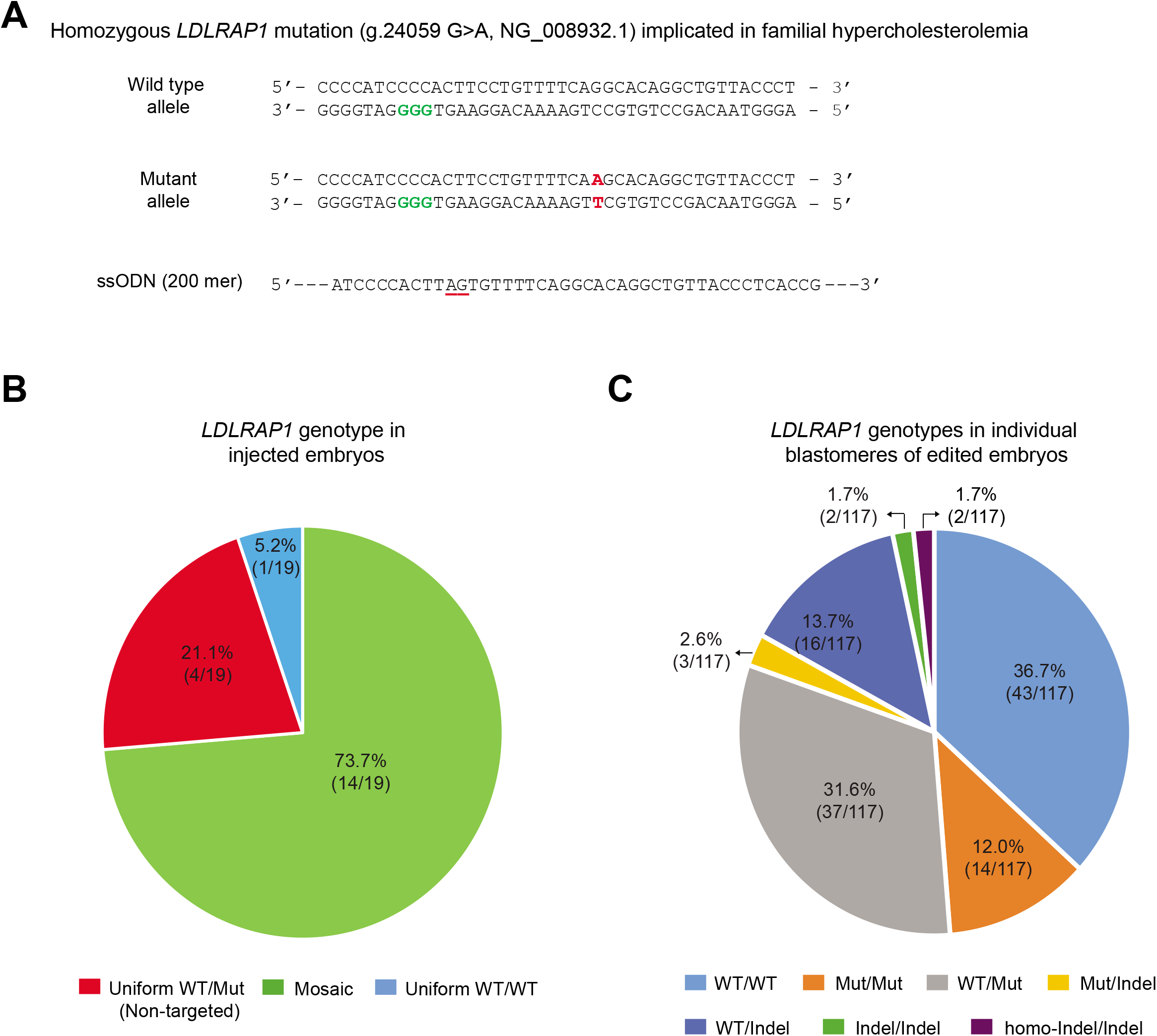
Gene conversion at the *LDLRAP1* locus. (**A**) DNA sequence of target *LDLRAP1* locus depicting WT (maternal) and mutant (paternal) alleles. DSBs were induced on both loci near the mutant g.24059 G>A locus (red fonts). Nucleotides shown in green indicate PAM, and red underlined nucleotides show substitutions in ssODN. (**B**) *LDLRAP1* genotypes in injected embryos. Note that all embryos were heterozygous *LDLRAP1^WT/Mut^* before injections. (**C**) Genotypes of individual blastomeres in injected embryos. DSBs are frequently repaired by gene conversion resulting in homozygous *LDLRAP1^WT/WT^, LDLRAP1^Mut/Mut^* or *LDLRAP1^homo-Indel/Indel^* genotypes.

The remaining 14 embryos were mosaic comprising a mixture of blastomeres with various genotypes (**Fig. 5B**). In 15 targeted embryos, 43/117 (36.7%) blastomeres lost the mutation and become homozygous *LDLRAP1^WT/WT^*. Conversely, 14/117 (12.0%) blastomeres lost the WT allele and become homozygous *LDLRAP1^Mut/Mut^* indicating reciprocal gene conversion using the mutant paternal allele as a template. In addition, 2/117 (1.7%) blastomeres were homozygous with identical indels (*LDLRAP1^homo-Indel/Indel^*). The remaining blastomeres (49.6%) were heterozygous carrying *LDLRAP1^WT/Mut^, LDLRAP1^WT/Indel^*, *LDLRAP1^Mut/Indel^* or *LDLRAP1^Indel/Indel^* genotypes (**Fig. 5C**). There was no evidence of HDR with ssODN.

These results validate our conclusions that DNA DSBs induced in human preimplantation embryos are frequently repaired by gene conversion. Based on observations of LOH at the heterozygous *LDLRAP1* locus (*A/G*), we estimate that the cumulative efficiency of gene conversion is more than 48.7%. In summary, based on targeting of *MYBPC3* and *LDLRAP1* loci, gene conversion occurs at high frequency (>40%) in human embryos even when targeting homozygous loci (**Fig. 6**). DSB repair by gene conversion often overlaps with NHEJ or HDR, resulting in LOH at the target locus. In contrast to heterozygous loci, induction of DSBs at homozygous loci may lead to HDR with exogenous DNA templates albeit at low efficiency. Thus, we conclude that gene conversion is one of the dominant DNA DSB repair pathways in human embryos.

**Fig 6.**
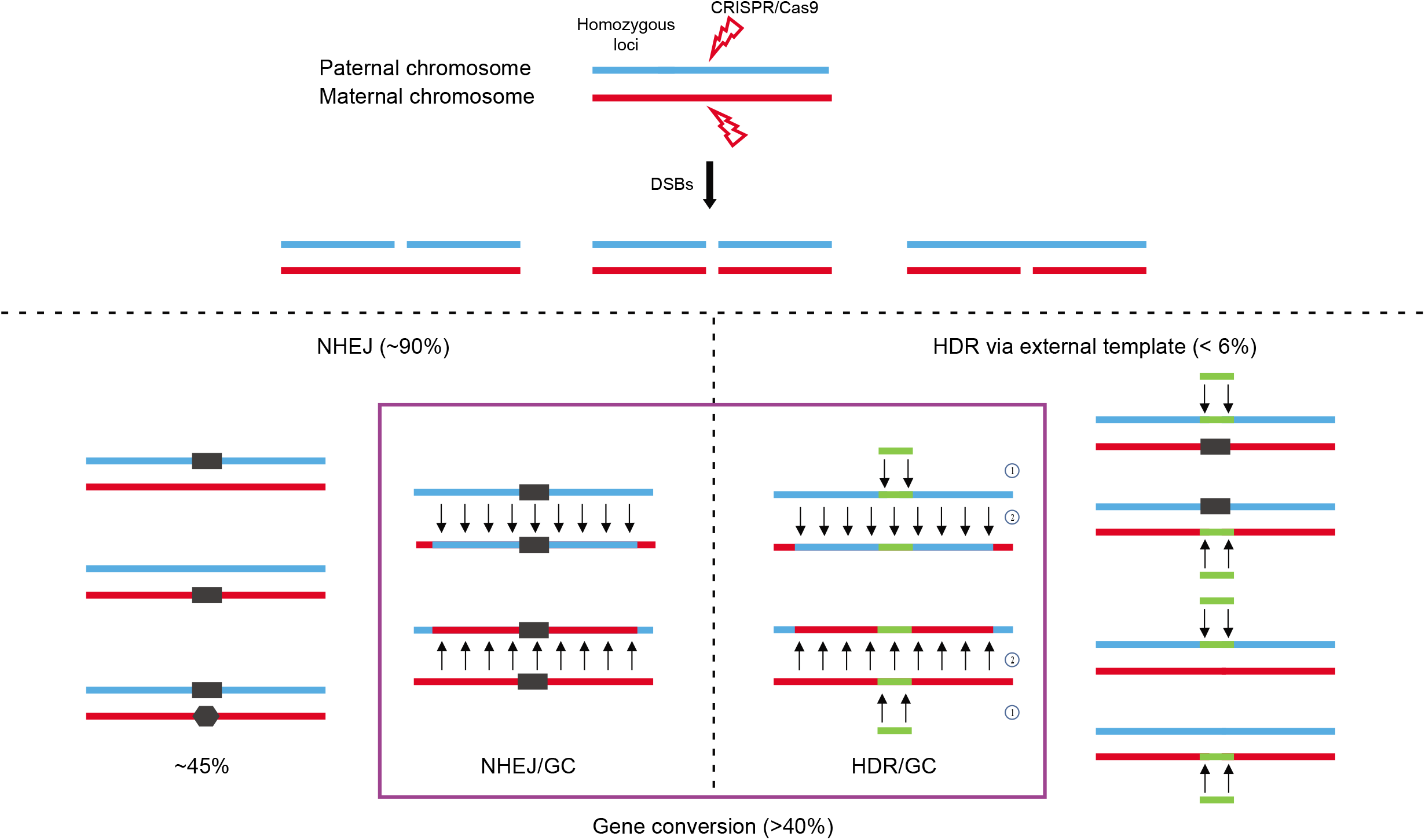
Summary of DNA DSB repair outcomes when targeting homozygous loci. Significant percentage of DSBs are repaired by NHEJ resulting in various indel mutations on one or both parental alleles. However, due to asynchronous targeting of two parental alleles, indel mutations on one chromosome can be copied to a homolog by gene conversion leading to LOH. In addition, HDR via exogenous DNA template was observed but at low frequency compared to that of NHEJ and gene conversion. Moreover, HDR via exogenous DNA template also can overlap with gene conversion, resulting in homozygous HDR and LOH.

## DISCUSSION

Meiotic and mitotic gene conversion was initially discovered in fungi as a phenomenon of non-Mendelian inheritance (15). In the past half century, gene conversion has been observed in different species from bacteria, plants, to mammals (16–18). In humans, gene conversion and associated LOH has been linked to inherited and acquired human diseases (9, 19–21). Conversely, the mutant alleles in some clinical cases involving heterozygous dominant mutations were reversed to the normal WT forms resulting in spontaneous gene therapy (22, 23).

Here, we demonstrate that DSBs in human preimplantation embryos are resolved primarily by NHEJ and gene conversion. These two DNA DSB repair mechanisms occur at similar frequencies and often compete within the same cell leading to the generation of identical indel mutations on both parental alleles. In contrast, the efficiency of HDR with exogenous DNA template was either completely absent when targeting heterozygous loci or found at low frequency when targeting homozygous loci.

We corroborate our previous findings that a significant portion of DSBs induced on a mutant paternal allele in heterozygous human embryos are repaired by gene conversion using an intact, WT maternal allele as a template. Interestingly, our results suggest that gene conversion is also active when targeting homozygous loci leading to the induction of homozygous indel mutations or homozygous HDR. We show that these novel homozygous loci are also associated with LOH at adjacent genomic regions due to erasure of one of the parental SNPs. Delayed access to one of the parental genomes in early human preimplantation embryos likely leads to selective induction of DSBs on one target locus while the second parental allele remains intact. In this case, the first DSB can be resolved by NHEJ resulting in a heterozygous indel mutation. Later, CRISPR/Cas9 locates and cuts the second intact allele, this DSB is resolved by gene conversion using the NHEJ-repaired homolog as a template.

Irrespective of whether CRIPSR-Cas9 was injected during fertilization or post-fertilization at the zygote stage, it appears that maternal genome is more readily accessible to CRISPR-Cas9 than the sperm DNA, leading to the preferential erasure of paternal SNPs in most embryos. One possible explanation is that during early post-fertilization stages, sperm chromatin is more tightly condensed and protected from nucleases by protamines compared to the oocyte genome (24).

Since induction of extensive homozygosity is a hallmark of gene conversion, sequencing-based validation of gene conversion outcomes is difficult. Indeed, our previous conclusions were challenged implying that the observed LOH can also be interpreted as complete loss of a parental allele due to large deletions (8, 25, 26). To address this issue, we incorporated here a FISH assay on ESCs derived from treated human embryos that provided visual, two signal confirmation that both alleles were intact. These results offer more definitive evidence of gene conversion in all tested samples. In addition, we screened all single blastomere DNA for large deletions using long-range PCR primers.

Given that DNA DSB repair by gene conversion is a conserved mechanism, it is likely a common outcome when programmable nucleases are introduced into embryos in other species. Indeed, an earlier mouse study suggested that selective induction of DSBs on the mutant *Crygc* locus in heterozygous mouse embryos can lead to gene conversion (27). The targeted allele was repaired using the WT homolog as template and more than 30% of live offspring lost the mutation and were healthy. In rats, allele-specific DSBs in heterozygous embryos were also repaired by an interallelic gene conversion at a frequency of 28% as judged by genetic and phenotypic analyses in live offspring (28).

It is likely that gene conversion outcomes remain largely undetected in most gene editing studies (29). Many animal studies reported cases of homozygous knock-out (identical indels) or knock-in (homozygous HDR) (30, 31). Based on our observations, some of these cases could be accounted for gene conversion. As discussed above, this can be only proven by detection of LOH in adjacent heterozygous loci. Mosaicism often masks gene conversion in pooled DNA samples, requiring single cell analysis.

Gene conversion could be applicable for future gene therapy to correct a mutant allele in heterozygous cells. To comply with strict requirements for germline gene therapy, gene conversion of heterozygous mutations back to the WT variants must be at much higher efficiency than observed in our present study. Conversely, the incidence of NHEJ must be significantly reduced. Extensive LOH in some blastomeres caused by long conversion tracts is a serious safety concern. LOH could lead to uncovering of preexisting heterozygous variants on a template genome leading to homozygosity of deleterious alleles and disease in offspring. Moreover, gene conversion may also erase parent-specific epigenetic DNA modifications leading to imprinting abnormalities.

Ultimately, HDR with exogenous DNA templates would be the most desirable and safe approach for germline gene therapy, especially for correcting homozygous mutations. However, the frequency of HDR in human embryos is currently low and unacceptable for therapeutic applications in human embryos.

## ACKNOWLEDGEMENTS

We thank the OHSU Institutional Review Board (IRB), innovative Research Advisory Panel (IRAP), Scientific Review Committee (SRC) and Data Safety Monitoring Committee (DSMC) for oversight and guidance of this study. We are grateful to all study participants for gamete and tissue donations and the Women’s Health Research Unit staff, IVF laboratory staff, University Fertility Consultants and the Reproductive Endocrinology and Infertility Division in the Department of Obstetrics and Gynecology, OHSU for their assistance in procurement of human gametes and Jun Wu and Juan Carlos Izpisua Belmonte for help with sgRNA selection. Studies conducted at the OHSU Center for Embryonic Cell and Gene Therapy were supported by OHSU institutional funds and a grant from the Burroughs Wellcome Fund. Studies conducted at Shandong university were supported by a grant from Chinese Thousand Talent Program. Studies conducted at Asan Medical Center were supported by a grant NRF-2015K1A4A3046807.

## Data and materials availability

All datasets generated during this study are included in this manuscript. Human ESC and iPSC lines are available to researchers upon OHSU IRB approval and signed OHSU MTA.

## Supplementary Information Text

### Study Oversight

Guidelines, policies and oversight defining research on human gametes and preimplantation embryos at Oregon Health & Science University (OHSU) were established by the Oregon Stem Cell and Embryo Research Oversight Committee (OSCRO). The study was approved by the OHSU Institutional Review Board (IRB) and included independent review by the OHSU Innovative Research Advisory Panel (IRAP) and OHSU Scientific Review Committee (SRC). The approved study was a subject for bi-annual external regulatory monitorings and Data Safety Monitoring Committee (DSMC) reviews.

### Informed Consent

Written informed consent was obtained from all subjects prior to enrollment in the study. Study subjects included sperm and egg donors and women with infertility undergoing IVF willing to donate their discarded immature oocytes for this study. Subjects were informed of risks to participation including risks associated with clinical procedures and loss of confidentiality.

### Study Participants

Adult sperm donors of 21-60 years of age carrying heritable *MYH7* or *LDLRAP1* mutations were identified and enrolled in this study by OHSU Knight Cardiovascular Institute physicians. In addition, healthy oocyte donors of 21-35 years of age were recruited locally, via print and webbased advertising.

### Compensation

Study participants providing gamete donations specifically for this research received financial compensation for their time, effort, and discomfort associated with the donation process at rates similar to gamete donation for fertility purposes. Infertility patients undergoing IVF whom donated immature oocytes did not receive any financial compensation.

### Ovarian Induction

Ovulation stimulation was managed by OHSU REI physicians as previously described (1) and followed established standards of care using a combination of self-administered injectable gonadotropins following 3-4 weeks ovarian suppression with combined oral contraceptives. Study participants self-administered medications for 8-12 days; the starting Follicle Stimulating Hormone (FSH) dose was 75–125 IU/day human Menopausal Gonadotropins (hMG) was adjusted per individual response using an established step-down regimen until the day of human Chorionic Gonadotropin (hCG) injection. GNRH antagonist was administered when the lead follicle was 14 mm in size. Subjects underwent ultrasound monitoring and blood draws for estradiol levels. hCG and/or Lupron was administered when two or more follicles measured >18 mm in diameter. Subjects underwent oocyte retrieval via transvaginal follicular aspiration 35 hours after hCG.

### Sperm Donation

Study subjects were provided an at home semen collection kit or collected their sample at OHSU REI clinic. Semen was washed, counted, and analyzed for volume, sperm count, motility, and morphology.

### Data and materials availability

OHSU IRB, Data Safety Monitoring Committee (DSMC) and Research Integrity control access to sequencing data and all samples generated during the course of this project. The sequence results will not be uploaded to a public database, no accession number. However, researchers my request access to this data by initiating a material transfer agreement with OHSU; approval may be granted after review by the DSMC, OHSU IRB, and Research Integrity.

## Supplementary materials and methods

### Skin fibroblast and peripheral blood mononuclear cell (PBMC) isolation and iPSCs derivation

A skin biopsy was collected from sperm donors, disaggregated into smaller pieces by incubation in collagenase IV for 30 minutes, and cells were then plated into 75-mm flasks in DMEM-F12 medium. Approximately, 10ml whole blood was collected into vacutainer tubes containing EDTA and 3ml was directly used for DNA extraction. The remaining blood was used for PBMC isolation with density gradient medium (Lymphoprep™, STEMCELL Technologies) and specialized tubes (SepMate™, STEMCELL Technologies), according to the manufacturer’s protocol. Skin fibroblasts or PBMC were treated with the CytoTune™-iPS 2.0 Sendai Reprogramming Kit (Life Technologies) to generate iPSCs, according to the manufacturer’s protocol.

### Human ESCs derivation

Zona pellucidae from blastocysts were removed with 0.5% protease (Sigma) and embryos were plated onto confluent feeder layers of mouse embryonic fibroblasts (mEF) and cultured for 6 days at 37°C, 3% CO_2_, 5% O_2_ and 92% N2 in ESC derivation medium. The medium consisted of DMEM/F12 (Invitrogen) with 0.1 mM nonessential amino acids, 1mM L-glutamine, 0.1mM β-mercaptoethanol, 5 ng/ml basic fibroblast growth factor, 10 μM ROCK inhibitor (Sigma), 10% fetal bovine serum (FBS) and 10% knockout serum replacement (KSR, Invitrogen). ESC colonies were manually dissociated and replated onto fresh mEFs for further propagation and analyses. FBS and ROCK inhibitor were omitted after the first passage of ESCs and KSR was increased to 20%.

### Fertilization and embryo culture

Mature MII oocytes were fertilized by intracytoplasmic sperm injection (ICSI) using fresh or frozen/thawed sperm as described earlier (2). Oocytes were placed into a 50μL droplet of HTF (modified human tubal fluid) medium supplemented with 10% HEPES, overlaid with mineral oil (Sage IVF, Cooper Surgical) and placed on the stage of an inverted microscope (Olympus IX71) equipped with a stage warmer and Narishige micromanipulators. A single sperm was drawn into ICSI micropipette and injected into the cytoplasm of each oocyte. Fertilized oocytes were then placed into dishes containing Global Medium (Life Global, IVF online) supplemented with 10% serum substitute supplement (Global 10% medium) and cultured at 37 °C in 6% CO_2_, 5% O_2_ and 89% N2 in an embryoscope time-lapse incubator (Vitrolife). Successful fertilization was determined approximately 18 hours after ICSI by noting the presence of two pronuclei and the second polar body extrusion.

### CRISPR/Cas9 design, selection and injection into human oocytes or zygotes

Multiple sgRNAs were designed for the *MYH7, MYBPC3* and *LDLRAP1* loci and synthesized by *in vitro* transcription using T7 polymerase (New England Biolabs) as described previously (1). Each sgRNA along with Cas9 and ssODN were transfected into blood or skin-derived iPSCs cells using Amaxa P3 Primary Cell 4D-Nucleofector Kit (Program). Three days after transfection, cells were harvested and DNA analyzed by targeted deep sequencing. CRISPR/Cas9 components that best performed in iPSCs were then selected for applications on human embryos. For the M-phase group, Cas9 protein (200ng/μl), sgRNA (100ng/μl) and ssODN (200ng/μl) were co-injected with sperm into the cytoplasm of each MII oocyte during ICSI procedure as described before (1). For the S-phase group, the CRISPR/Cas9 components were injected into cytoplasm of pronuclear stage zygotes 18 hours after ICSI.

### Blastomere isolation and whole genome DNA amplification

Injected oocytes or zygotes were cultured to the 4-8 cell stage and used for single blastomere analyzes as described (1). Briefly, the zona pellucida of cleaving embryos was removed using acid Tyrode’s solution (NaCl 8 mg/ml, KCl 0.2 mg/ml, CaCl_2_.2H_2_O 2.4 mg/ml, MgCl_2_.6H_2_O 0.1 mg/ml, glucose 1 mg/ml, PVP 0.04 mg/ml). Embryos were then briefly exposed in a trypsin solution, and individual blastomeres were mechanically separated using a micromanipulation pipette. Each blastomere was then placed into 0.2 ml PCR tube containing 4 μl PBS and stored at −80°C. Whole genome amplification was performed using a REPLI-g Single Cell Kit (Qiagen), according to the manufacturer’s protocol. Briefly, frozen/thawed tubes containing blastomeres were treated with denaturation solution mix and incubated at 65°C for 10 min. A master mix containing buffer and DNA polymerase was then added to each tube. The amplification reaction processed for 8 hours at 30°C in a PCR thermocycler. Whole genome amplification product was then diluted 100 times and used for downstream applications.

### Genotyping, Sanger sequencing and Long-range PCR

The target region for MYH7, MYBPC3 and LDLRAP1 loci and SNP sites were amplified with PCR primers using PCR platinum SuperMix High Fidelity Kit (Life Technologies). PCR products were purified by ExoSAP-IT reagent (Affymetrix), single purify condition were: 5ul PCR product with 2ul of ExoSAP-IT reagent, 37oC for 15 min then 80oC for 15min. Then purified PCR product were sequenced by Sanger and analyzed by SnapGene^®^ Viewer. Long-range PCR amplifications were performed by using TaKaRa LA Taq DNA Polymerase (Clontech) as described previously (3).

### Next Generation Sequencing and Bioinformatics Analysis

Genomic DNA from the gamete donors was processed by WES. Sequencing libraries were prepared according to the manufacturer’s instruction for BGISEQ WES library preparation. DNA fragments were hybridized to the exome array BGI-V4 chip for enrichment, and high-throughput sequencing was performed for each captured library on BGISEQ-500 sequencing platform with paired-end 100 bp (PE100) strategy at an average depth of 69.12X and coverage of 97.97% on the target region.

All sequencing data were first processed by filtering adaptor and removing low quality reads or reads with high percentage of N bases using SOAPnuke software v1.5.2. The data were then aligned to the human genome assembly hg19 (GRCh37) using Burrows-Wheeler Aligner (BWA) software v0.7.15. To ensure accurate variant calling, best practices for variant analysis with the Genome Analysis Toolkit (GATK) were followed. Local realignment around InDels and base quality score recalibration were performed using GATK, with duplicate reads removed by Picard v2.5.0 tools (https://broadinstitute.github.io/picard/). All genomic variations, including SNPs were detected by HaplotypeCaller tool in GATK. The hard-filtering method was subsequently applied to achieve high-confident variant calls. Thereafter, the SnpEff tool (http://snpeff.sourceforge.net/SnpEff_manual.html) was applied to perform a series of annotations for variants.

### DNA fluorescence in situ hybridization (FISH)

FISH analyses were carried on metaphase arrested ESCs as previously described (4). Briefly, ESCs were treated with KaryoMAX Colcemide (Life Technologies) at a final concentration of 200 ng/mL for 1.5 hours at 37°C. Treated cells were then detached by 0.25% trypsin/EDTA and incubated in hypotonic 0.075M KCL for 20 minutes. Cells were next fixed with methanol: acetic acid (3:1 v/v) and dropped onto a slide and dried on a hot plate at 60°C. The samples were dehydrated using ethanol (70%, 85%, and 100%) for 1 minute in each and dried in air. FISH probes specific for *MYBPC3* (11p11.2 locus) were labeled using Green-dUTP and the *MYH7* (14q11.2 locus) was labeled using Red-dUTP (Empire Genomics). Slides were applied with the probe mixture, covered with an 18mm^2^ coverslip, and incubated in a humidified Thermobrite^®^ system (Leica) set at 73°C for 2 minutes, and then 37°C for 16 hours. The incubated slides were rinsed with washing solution 1 (0.3% Igepal/0.4xSSC) and washing solution 2 (0.1% Igepal/2xSSC). Slides were mounted in ProLong™ Gold Antifade Mountant with DAPI (Life Technologies) and observed using a fluorescence microscopy equipped with a cooled CCD camera. Images were captured and analyzed by ISIS analysis software (MetaSystem GmbH).

**Fig. S1.**
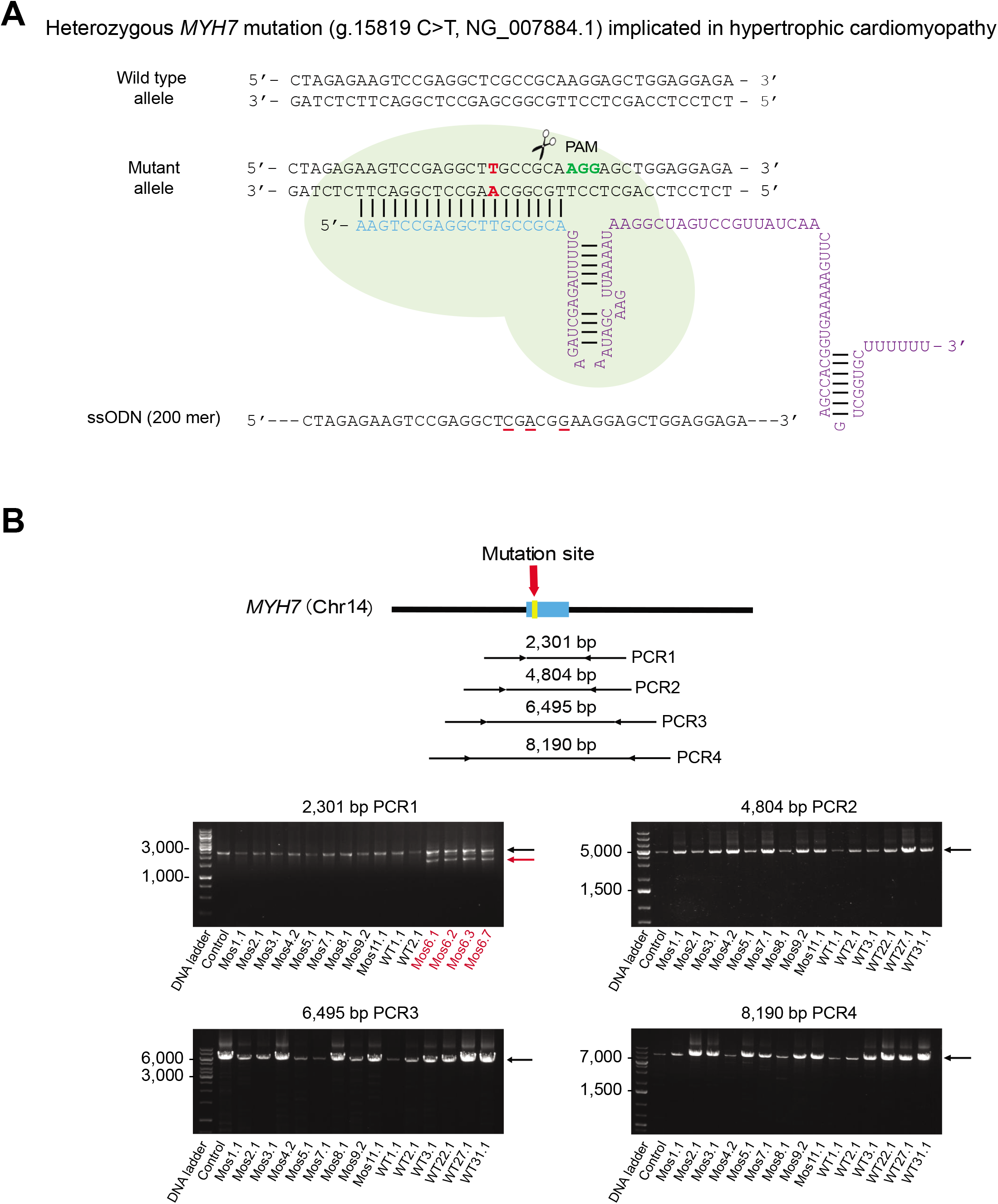
*MYH7* target region and long-range PCR for detection of large deletions. **(A)** Human wildtype and mutant *MYH7* g.15819 C>T locus implicated in hypertrophic cardiomyopathy. sgRNA was designed to target the mutant allele. Red font nucleotides show mutation, green indicate PAM, and red underlined nucleotides show substitutions in ssODN. **(B)** Long-range PCR electrophoresis screening for large deletions. DNA from each individual blastomere was analyzed using 4 pairs of long-range PCR primers spanning the *MYH7* g.15819 *C>T* locus. Agarose gel electrophoregram showing 2301 bp (PCR1), 4804 bp (PCR2), 6495 bp (PCR3) and 8190 bp (PCR4) target fragments in blastomeres from mosaic and uniform WT embryos. All blastomeres designated as *MYH7^WT/WT^* did not show any secondary smaller size bands indicating lack of deletions. Four blastomeres (shown in red fonts) from one embryo with *MYH7^WT/indel^* genotype carried a large deletion of 652 bp detectable by electrophoresis (pointed by red arrow). Black arrows show expected size PCR brands.

**Fig. S2.**
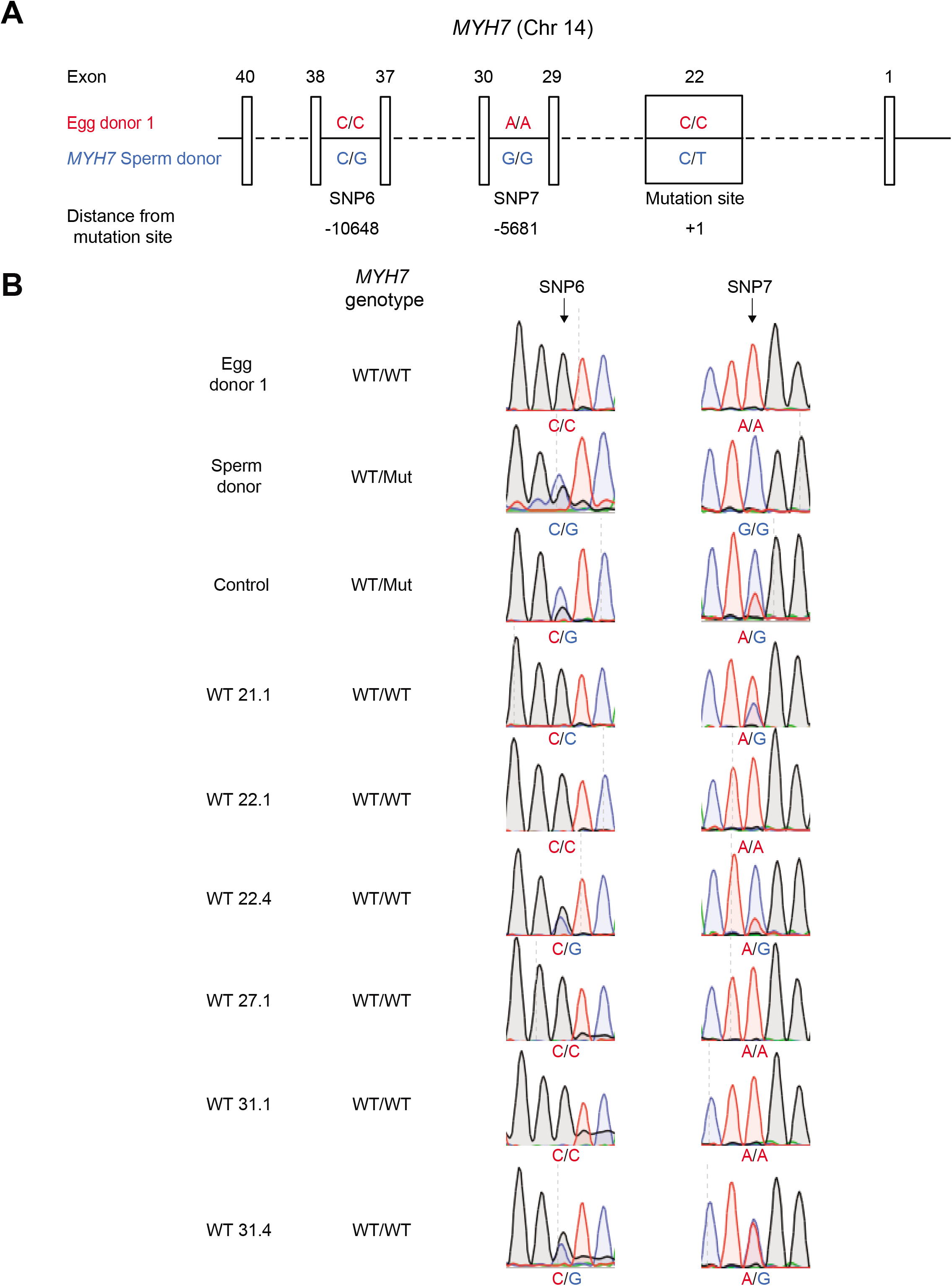
LOH at adjacent heterozygous loci due to gene conversion in *MYH7* targeted human embryos. (A) A schematic sequence map of two informative heterozygous loci (SNP6 and 7) upstream of MYH7 mutant locus in embryos generated from egg donor 1 and MYH7 sperm donor. Red font nucleotides represent maternal origin, blue indicate paternal and boxes indicate exons. (B) Sanger sequencing chromatographs of SNP6 and 7 loci in individual blastomeres from injected embryos. Note that some MYH7WT/WT blastomeres lost heterozygosity at both SNP6 and SNP7 loci indicative of gene conversion. See more details in **Table 1**.

**Fig. S3.**
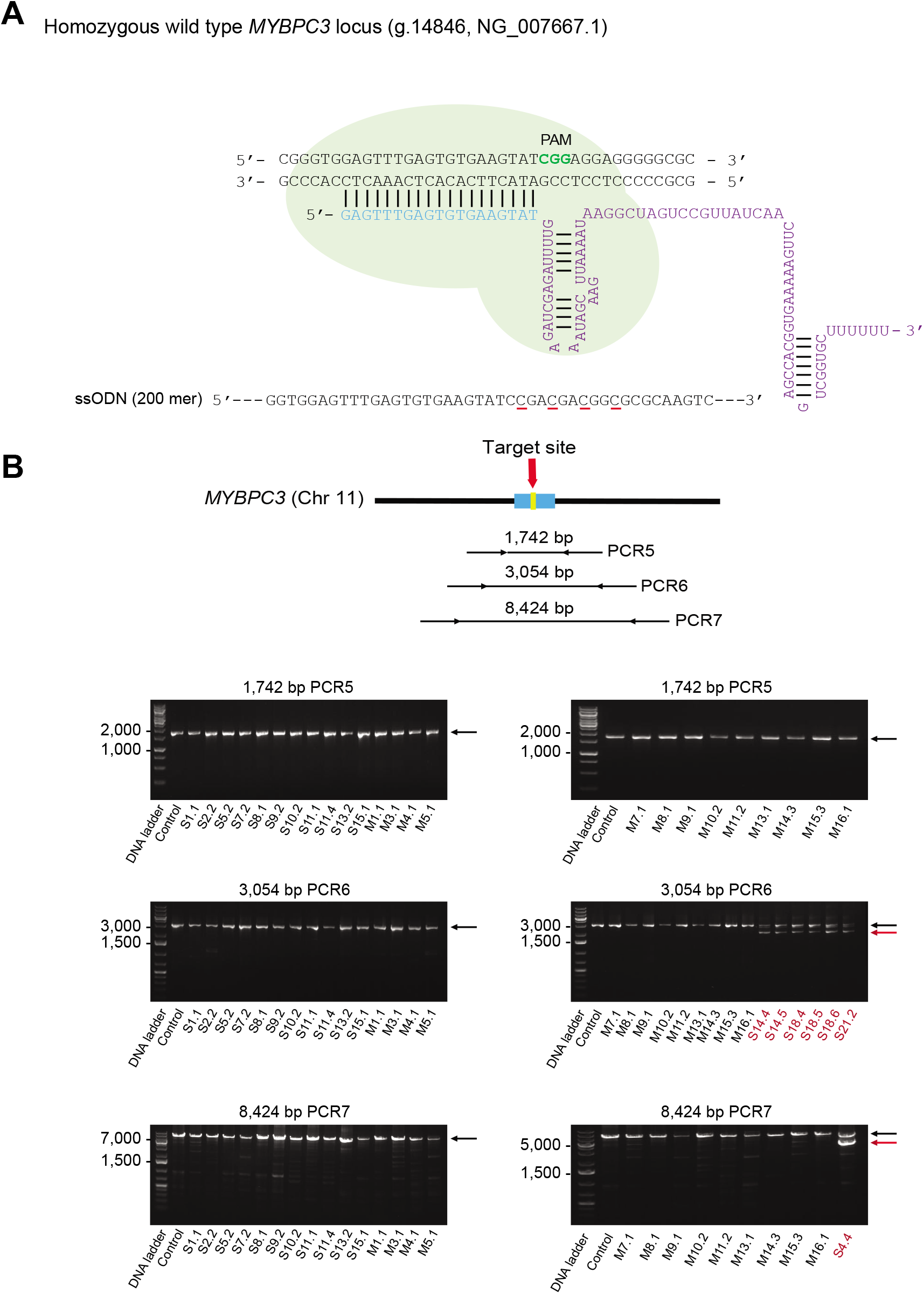
*MYBPC3* target region and long-range PCR for detection of large deletions. **(A)** Human wildtype *MYBPC3* target locus. Note that a 4 bp deletion in this locus is associated with hypertrophic cardiomyopathy (1). Pre-selected sgRNA targets both wild type alleles. Green font nucleotides indicate PAM, underlined nucleotides show substitutions in ssODN. **(B)** Long-range PCR electrophoresis screening for large deletions in *MYBPC3* region. DNA from individual blastomeres was analyzed using 3 pairs of long-range PCR primers spanning the target *MYH7* locus. Agarose gel electrophoregrams showing bands of expected size of 1742 bp (PCR5), 3054 bp (PCR6) and 8424 bp (PCR7) in all *MYBPC3^homo-Indel/Indel^, MYBPC3^HDR/HDR^* (S11.4) and control blastomeres (shown by black fonts). Note that 6 blastomeres (shown in red fonts) with *MYBPC3^Indel/Indel^* genotype showed a secondary band indicating large deletions (gel PCR 6, pointed by red arrows). Likewise, one blastomere (S4.4, shown by red fonts in PCR7 gel) with *MYBPC3^Indel/Indel^* genotype had largest deletion (pointed by red arrow). Black arrows show expected size PCR brands.

**Table S1.**
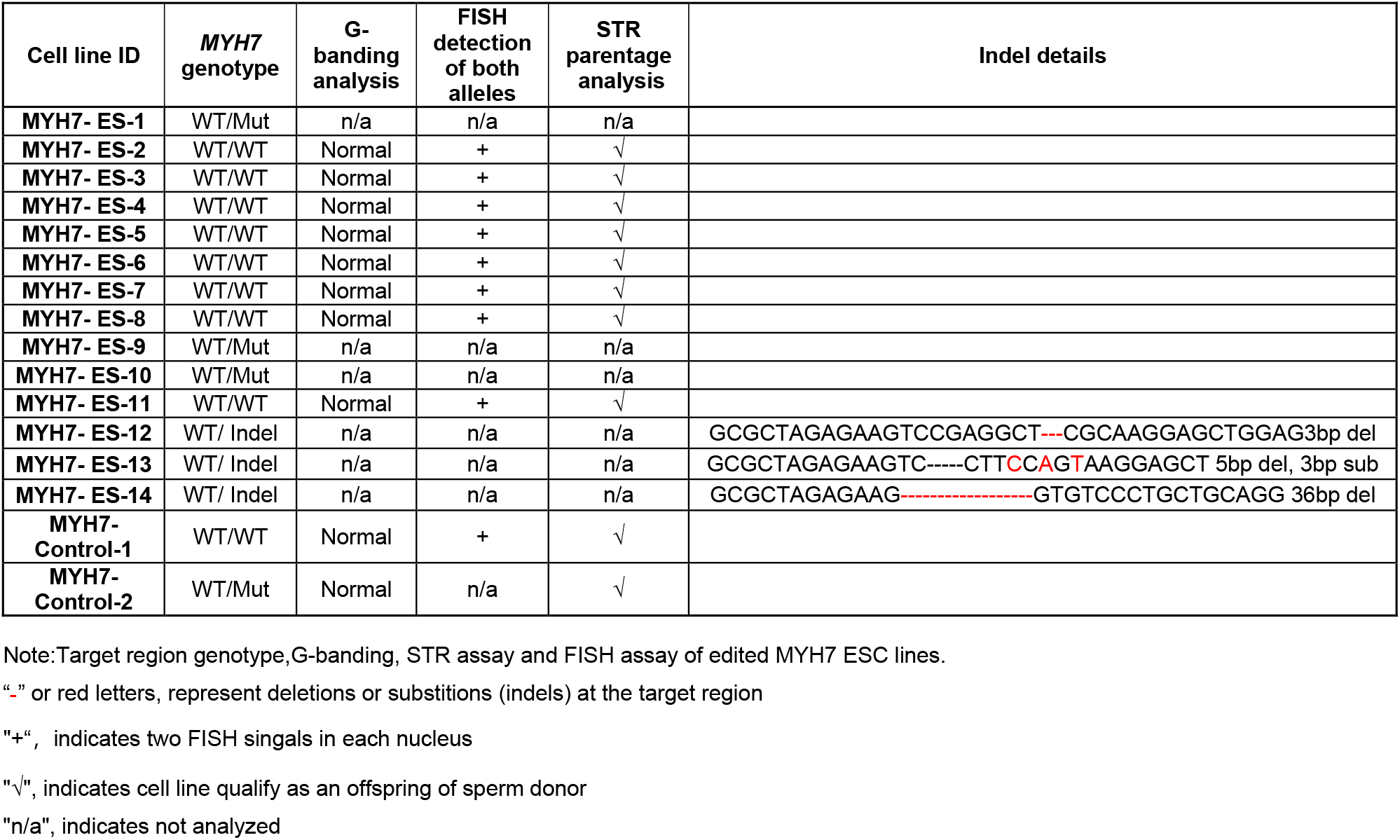
*MYH7* ESC lines derived from CRISPR/Cas9 injected and control blastocysts

**Table S2.**
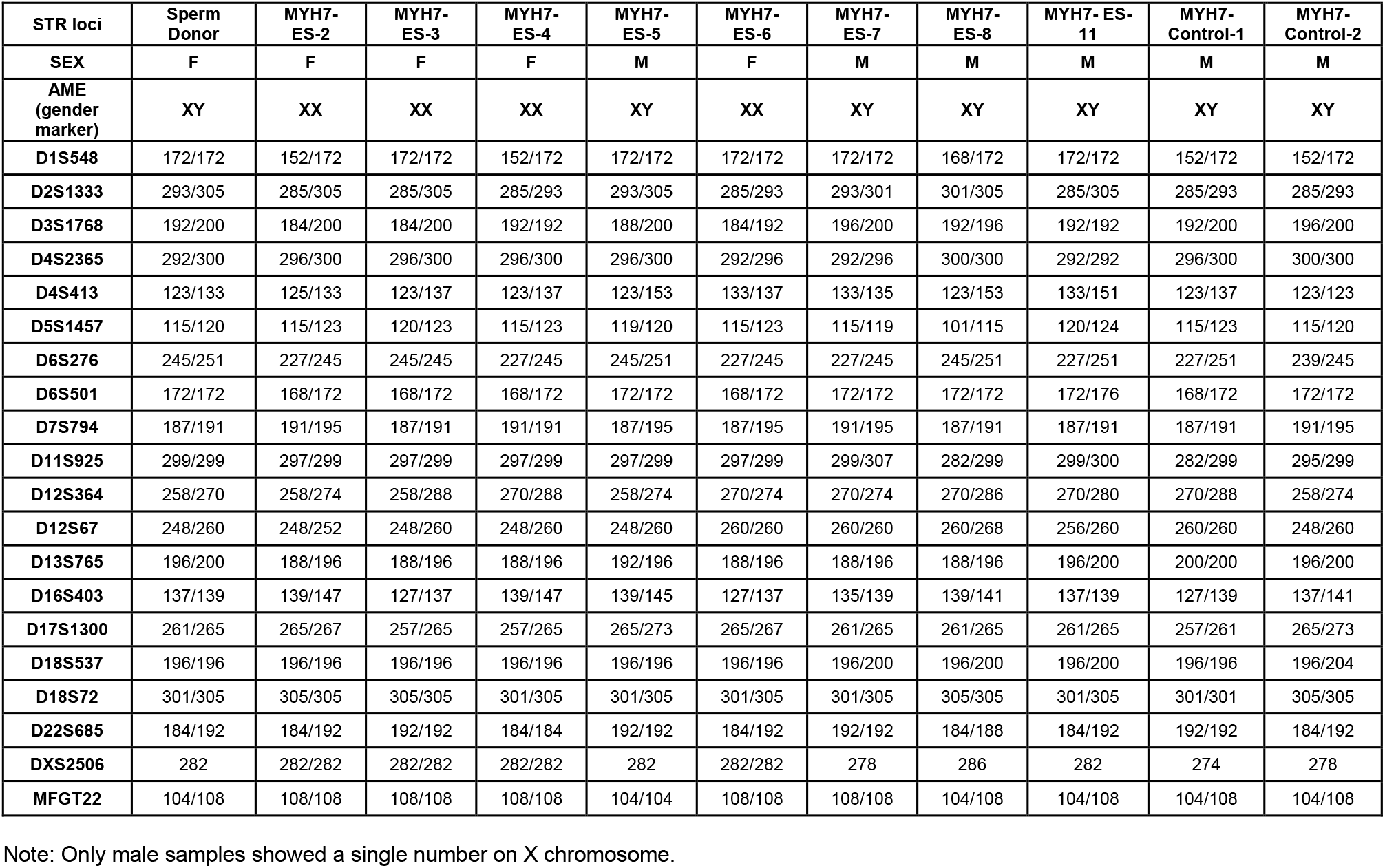
STR genotypes of injected *MYH7* WT/WT ESC lines

| STR loci | Sperm Donor | MYH7-ES-2 | MYH7-ES-3 | MYH7-ES-4 | MYH7-ES-5 | MYH7-ES-6 | MYH7-ES-7 | MYH7-ES-8 | MYH7-ES-11 | MYH7-Control-1 | MYH7-Control-2 |
| --- | --- | --- | --- | --- | --- | --- | --- | --- | --- | --- | --- |
| SEX | F | F | F | F | M | F | M | M | M | M | M |
| AME (gender marker) | XY | XX | XX | XX | XY | XX | XY | XY | XY | XY | XY |
| D1S548 | 172/172 | 152/172 | 172/172 | 152/172 | 172/172 | 172/172 | 172/172 | 168/172 | 172/172 | 152/172 | 152/172 |
| D2S1333 | 293/305 | 285/305 | 285/305 | 285/293 | 293/305 | 285/293 | 293/301 | 301/305 | 285/305 | 285/293 | 285/293 |
| D3S1768 | 192/200 | 184/200 | 184/200 | 192/192 | 188/200 | 184/192 | 196/200 | 192/196 | 192/192 | 192/200 | 196/200 |
| D4S2365 | 292/300 | 296/300 | 296/300 | 296/300 | 296/300 | 292/296 | 292/296 | 300/300 | 292/292 | 296/300 | 300/300 |
| D4S413 | 123/133 | 125/133 | 123/137 | 123/137 | 123/153 | 133/137 | 133/135 | 123/153 | 133/151 | 123/137 | 123/123 |
| D5S1457 | 115/120 | 115/123 | 120/123 | 115/123 | 119/120 | 115/123 | 115/119 | 101/115 | 120/124 | 115/123 | 115/120 |
| D6S276 | 245/251 | 227/245 | 245/245 | 227/245 | 245/251 | 227/245 | 227/245 | 245/251 | 227/251 | 227/251 | 239/245 |
| D6S501 | 172/172 | 168/172 | 168/172 | 168/172 | 172/172 | 168/172 | 172/172 | 172/172 | 172/176 | 168/172 | 172/172 |
| D7S794 | 187/191 | 191/195 | 187/191 | 191/191 | 187/195 | 187/195 | 191/195 | 187/191 | 187/191 | 187/191 | 191/195 |
| D11S925 | 299/299 | 297/299 | 297/299 | 297/299 | 297/299 | 297/299 | 299/307 | 282/299 | 299/300 | 282/299 | 295/299 |
| D12S364 | 258/270 | 258/274 | 258/288 | 270/288 | 258/274 | 270/274 | 270/274 | 270/286 | 270/280 | 270/288 | 258/274 |
| D12S67 | 248/260 | 248/252 | 248/260 | 248/260 | 248/260 | 260/260 | 260/260 | 260/268 | 256/260 | 260/260 | 248/260 |
| D13S765 | 196/200 | 188/196 | 188/196 | 188/196 | 192/196 | 188/196 | 188/196 | 188/196 | 196/200 | 200/200 | 196/200 |
| D16S403 | 137/139 | 139/147 | 127/137 | 139/147 | 139/145 | 127/137 | 135/139 | 139/141 | 137/139 | 127/139 | 137/141 |
| D17S1300 | 261/265 | 265/267 | 257/265 | 257/265 | 265/273 | 265/267 | 261/265 | 261/265 | 261/265 | 257/261 | 265/273 |
| D18S537 | 196/196 | 196/196 | 196/196 | 196/196 | 196/196 | 196/196 | 196/200 | 196/200 | 196/200 | 196/196 | 196/204 |
| D18S72 | 301/305 | 305/305 | 305/305 | 301/305 | 301/305 | 301/305 | 301/305 | 305/305 | 301/305 | 301/301 | 305/305 |
| D22S685 | 184/192 | 184/192 | 192/192 | 184/184 | 192/192 | 184/192 | 192/192 | 184/188 | 184/192 | 192/192 | 184/192 |
| DXS2506 | 282 | 282/282 | 282/282 | 282/282 | 282 | 282/282 | 278 | 286 | 282 | 274 | 278 |
| MFGT22 | 104/108 | 108/108 | 108/108 | 108/108 | 104/104 | 108/108 | 108/108 | 104/108 | 104/108 | 104/108 | 104/108 |

Note: Only male samples showed a single number on X chromosome.

**Table S3.** SNP genotypes in mosaic embryos from egg donor 1 and *MYH7* mutant sperm donor

| Embryo ID | Blastomere ID | *MYH7* on-target genotype | SNP1 (rs7493807, ~ -3424 kb) | SNP2 (rs12896494, -77054 bp) | SNP3 (rs723840, -49155 bp) | SNP4 (rs8006357, -40423 bp) | SNP5 (rs2277474, -19529 bp) | SNP6 (rs41285540, -10648 bp) | SNP7 (rs3729820, -5681 bp) | SNP8 (rs7157716, -1164 bp) | SNP9 (rs1951154, +3415 bp) | SNP10 (rs2069542, +6743 bp) | SNP11 (rs2069540, +8702 bp) | SNP12 (rs2295705, +45381 bp) | SNP13 (rs55633823, ~+82060 kb) |
| --- | --- | --- | --- | --- | --- | --- | --- | --- | --- | --- | --- | --- | --- | --- | --- |
| Egg donor 1 | n/a | WT/WT | A | T | C | T | C | C | A | A | A | G | G | G | T |
| MYH7 sperm donor | n/a | WT/Mut | G | C | C/T | T/C | C/T | C/G | G | A/G | A/G | G/A | G/A | A | C |
| Control 12 | 12.1 | WT/Mut | A/G | C/T | C/T | T/C | C/T | C/G | A/G | A/G | A/G | G/A | G/A | G/A | T/C |
| MYH7-Mos13 | 13.1 | WT/WT | A | T | C | T | C | C | A | A | A | G | G | G | n/a |
|  | 13.2 | WT/WT | A/G | C/T | C/T | T/C | C/T | C/G | A/G | A/G | A | G | G | G | T |
|  | 13.3 | WT/WT | A/G | C/T | C/T | T/C | C/T | C/G | A/G | A | A/G | G/A | G/A | G/A | n/a |
|  | 13.5 | WT/WT | A | T | C | T | C | C | A | A | A/G | G/A | G/A | G/A | n/a |
|  | 13.6 | WT/Indel | A/G | C/T | C/T | T/C | C/T | C/G | A/G | A/G | A/G | G/A | G/A | G/A | T/C |
| MYH7-Mos14 | 14.2 | WT/WT | A/G | C/T | C/T | T/C | C/T | C/G | A/G | A/G | A/G | G/A | G/A | G/A | n/a |
|  | 14.3 | WT/WT | A/G | C/T | C | T | C | C | A | A | A | G | G | G | T |
|  | 14.6 | WT/WT | A/G | C/T | C/T | T/C | T | G | G | G | A | G/A | G/A | G/A | n/a |
|  | 14.7 | WT/WT | A/G | C/T | C/T | T/C | C/T | C/G | A/G | A | A | G | G | G | T |
|  | 14.4 | WT/Indel | A/G | C/T | C/T | T/C | C/T | C/G | A/G | A/G | A/G | G/A | G/A | G/A | T/C |
| MYH7-Mos15 | 15.1 | WT/WT | A/G | C/T | C/T | T/C | C/T | C/G | A/G | A/G | A | G | G | G | T |
|  | 15.2 | WT/WT | A | T | C | T | C | C | A | A | A | G | G | G | T |
|  | 15.4 | WT/WT | A/G | C/T | C/T | T/C | C | C | A | A | A | G/A | G/A | G/A | T/C |
|  | 15.5 | WT/WT | A | T | C | T | C | C | A | A | A | G | G | G | T |
|  | 15.6 | WT/WT | A | T | C | T | C | C | A | A | A | G | G | G | n/a |
|  | 15.7 | WT/WT | A/G | C/T | C | T | C | C | A | A | A | G | G | G | T |
|  | 15.3 | WT/Indel | A/G | C/T | C/T | T/C | C/T | C/G | A/G | T/C | A/G | G/A | G/A | G/A | T/C |

**Table S4.**
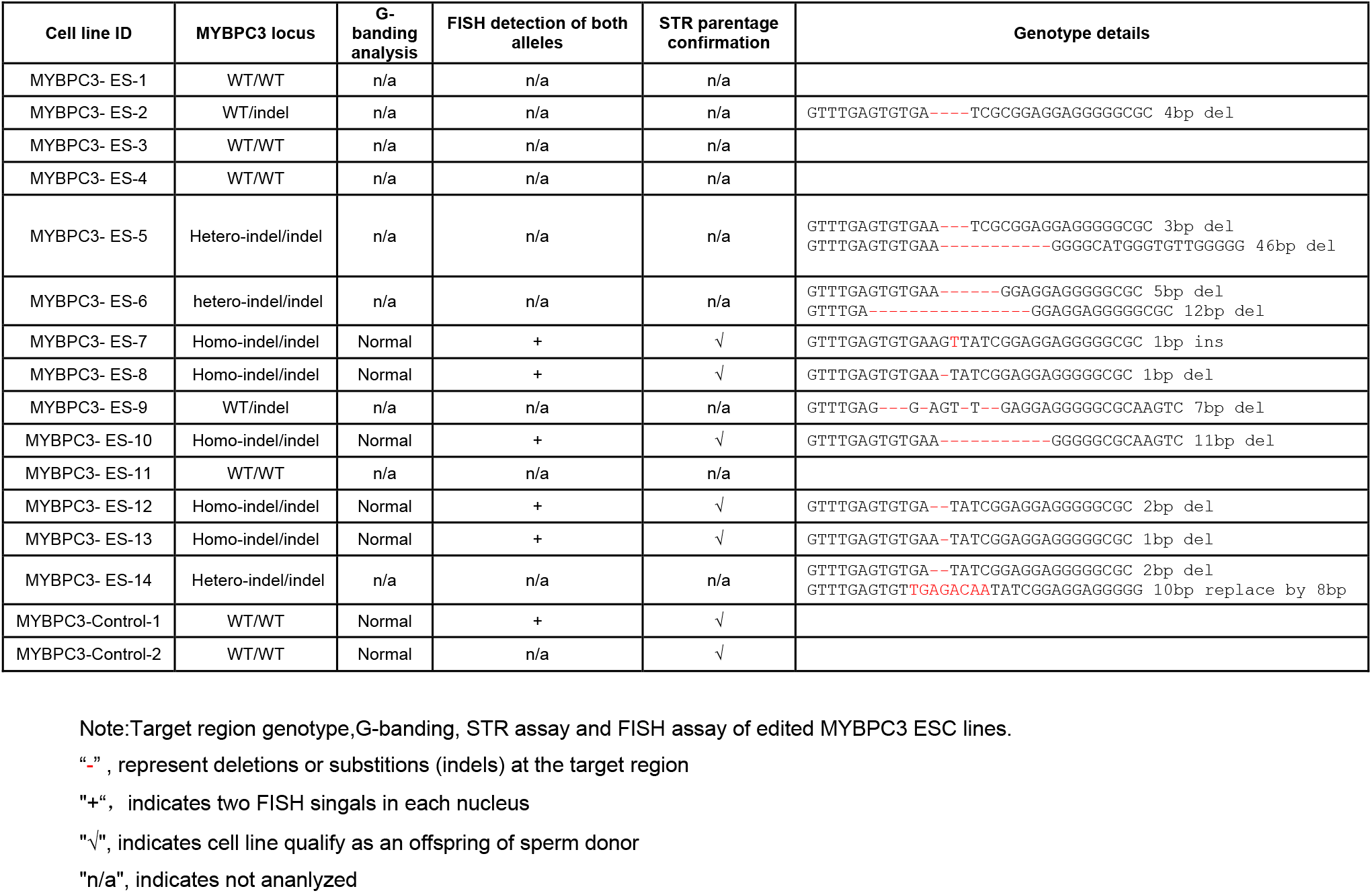
*MYBPC3* ESC lines derived from injected and control embryos

**Table S5.**
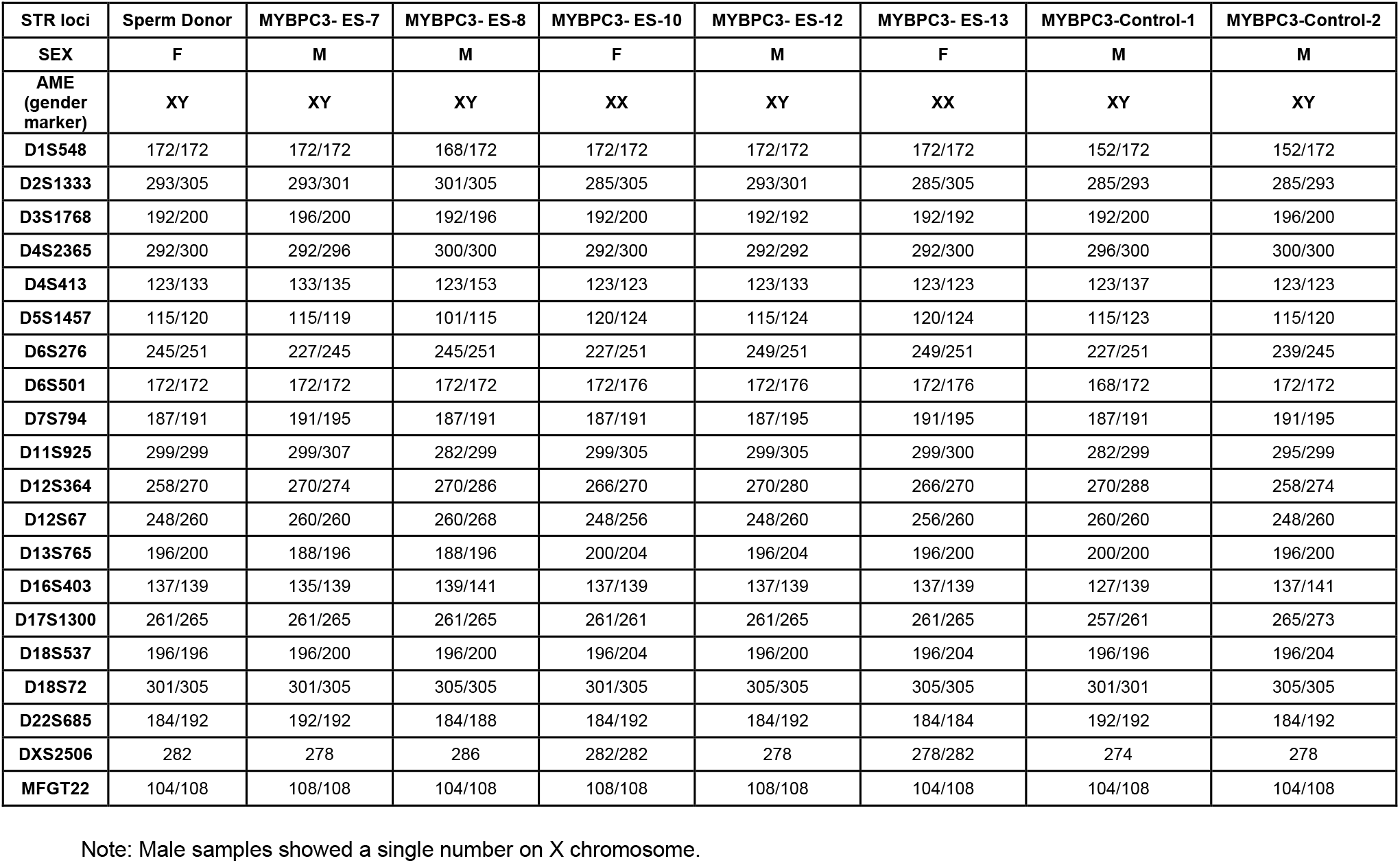
STR genotypes in edited *MYBPC3* homo-Indel/Indel ESC lines

| STR loci | Sperm Donor | MYBPC3- ES-7 | MYBPC3- ES-8 | MYBPC3- ES-10 | MYBPC3- ES-12 | MYBPC3- ES-13 | MYBPC3-Control-1 | MYBPC3-Control-2 |
| --- | --- | --- | --- | --- | --- | --- | --- | --- |
| SEX | F | M | M | F | M | F | M | M |
| AME<br>(gender<br>marker) | XY | XY | XY | XX | XY | XX | XY | XY |
| D1S548 | 172/172 | 172/172 | 168/172 | 172/172 | 172/172 | 172/172 | 152/172 | 152/172 |
| D2S1333 | 293/305 | 293/301 | 301/305 | 285/305 | 293/301 | 285/305 | 285/293 | 285/293 |
| D3S1768 | 192/200 | 196/200 | 192/196 | 192/200 | 192/192 | 192/192 | 192/200 | 196/200 |
| D4S2365 | 292/300 | 292/296 | 300/300 | 292/300 | 292/292 | 292/300 | 296/300 | 300/300 |
| D4S413 | 123/133 | 133/135 | 123/153 | 123/123 | 123/133 | 123/123 | 123/137 | 123/123 |
| D5S1457 | 115/120 | 115/119 | 101/115 | 120/124 | 115/124 | 120/124 | 115/123 | 115/120 |
| D6S276 | 245/251 | 227/245 | 245/251 | 227/251 | 249/251 | 249/251 | 227/251 | 239/245 |
| D6S501 | 172/172 | 172/172 | 172/172 | 172/176 | 172/176 | 172/176 | 168/172 | 172/172 |
| D7S794 | 187/191 | 191/195 | 187/191 | 187/191 | 187/195 | 191/195 | 187/191 | 191/195 |
| D11S925 | 299/299 | 299/307 | 282/299 | 299/305 | 299/305 | 299/300 | 282/299 | 295/299 |
| D12S364 | 258/270 | 270/274 | 270/286 | 266/270 | 270/280 | 266/270 | 270/288 | 258/274 |
| D12S67 | 248/260 | 260/260 | 260/268 | 248/256 | 248/260 | 256/260 | 260/260 | 248/260 |
| D13S765 | 196/200 | 188/196 | 188/196 | 200/204 | 196/204 | 196/200 | 200/200 | 196/200 |
| D16S403 | 137/139 | 135/139 | 139/141 | 137/139 | 137/139 | 137/139 | 127/139 | 137/141 |
| D17S1300 | 261/265 | 261/265 | 261/265 | 261/261 | 261/265 | 261/265 | 257/261 | 265/273 |
| D18S537 | 196/196 | 196/200 | 196/200 | 196/204 | 196/200 | 196/204 | 196/196 | 196/204 |
| D18S72 | 301/305 | 301/305 | 305/305 | 301/305 | 305/305 | 305/305 | 301/301 | 305/305 |
| D22S685 | 184/192 | 192/192 | 184/188 | 184/192 | 184/192 | 184/184 | 192/192 | 184/192 |
| DXS2506 | 282 | 278 | 286 | 282/282 | 278 | 278/282 | 274 | 278 |
| MFGT22 | 104/108 | 108/108 | 104/108 | 108/108 | 108/108 | 104/108 | 104/108 | 104/108 |

Note: Male samples showed a single number on X chromosome.

**Table S6.** LOH in homo-Indel/indel blastomeres from *MYBPC3* edited emrbyos (S-phase)

| Sample ID | *MYBPC3* on-target genotype | SNP14 genotype<br>(rs10769253, -2085 bp) |
| --- | --- | --- |
| Egg donor 1 | WT/WT | G/G |
| Sperm donor | WT/WT | A/A |
| MYBPC3-S8.1 | homo-Indel/Indel | G/G |
| MYBPC3-S8.2 | homo-Indel/Indel | G/G |
| MYBPC3-S8.3 | homo-Indel/Indel | A/A |
| MYBPC3-S9.1 | homo-Indel/Indel | G/G |
| MYBPC3-S9.2 | homo-Indel/Indel | A/A |
| MYBPC3-S9.3 | homo-Indel/Indel | A/A |
| MYBPC3-S9.6 | homo-Indel/Indel | G/A |
| MYBPC3-S9.7 | homo-Indel/Indel | G/A |
| MYBPC3-S10.2 | homo-Indel/Indel | G/A |
| MYBPC3-S10.4 | homo-Indel/Indel | A/A |
| MYBPC3-S10.5 | homo-Indel/Indel | A/A |
| MYBPC3-S10.6 | homo-Indel/Indel | G/A |
| MYBPC3-S10.8 | homo-Indel/Indel | A/A |
| MYBPC3-S10.9 | homo-Indel/Indel | A/A |
| MYBPC3-S11.1 | homo-Indel/Indel | A/A |
| MYBPC3-S11.3 | homo-Indel/Indel | G/G |
| MYBPC3-S12.5 | homo-Indel/Indel | G/A |
| MYBPC3-S13.1 | homo-Indel/Indel | G/G |
| MYBPC3-S13.2 | homo-Indel/Indel | A/A |
| MYBPC3-S13.3 | homo-Indel/Indel | G/G |
| MYBPC3-S13.4 | homo-Indel/Indel | G/G |
| MYBPC3-S13.5 | homo-Indel/Indel | G/G |
| MYBPC3-S15.1 | homo-Indel/Indel | A/A |
| MYBPC3-S15.2 | homo-Indel/Indel | A/A |
| MYBPC3-S16.1 | homo-Indel/Indel | G/G |
| MYBPC3-S16.3 | homo-Indel/Indel | G/G |
| MYBPC3-S16.4 | homo-Indel/Indel | G/A |
| MYBPC3-S18.1 | homo-Indel/Indel | G/G |
| MYBPC3-S18.3 | homo-Indel/Indel | G/G |
| MYBPC3-S19.1 | homo-Indel/Indel | G/G |
| MYBPC3-S19.2 | homo-Indel/Indel | G/A |
| MYBPC3-S19.3 | homo-Indel/Indel | G/G |
| MYBPC3-S19.4 | homo-Indel/Indel | G/A |
| MYBPC3-S19.5 | homo-Indel/Indel | G/A |
| MYBPC3-S19.6 | homo-Indel/Indel | G/G |
| MYBPC3-S19.7 | homo-Indel/Indel | G/G |
| MYBPC3-S19.8 | homo-Indel/Indel | G/G |
| MYBPC3-S19.9 | homo-Indel/Indel | G/G |
| MYBPC3-S19.10 | homo-Indel/Indel | G/G |
| MYBPC3-S20.1 | homo-Indel/Indel | G/A |
| MYBPC3-S20.2 | homo-Indel/Indel | A/A |
| MYBPC3-S20.3 | homo-Indel/Indel | A/A |
| MYBPC3-S20.4 | homo-Indel/Indel | A/A |
| MYBPC3-S21.1 | homo-Indel/Indel | G/G |
| MYBPC3-S21.4 | homo-Indel/Indel | G/G |
| MYBPC3-S21.6 | homo-Indel/Indel | G/G |
| MYBPC3-S21.8 | homo-Indel/Indel | G/G |

**Table S7.** LOH in homo-Indel/indel blastomeres from MYBPC3 edited emrbyos (M-phase)

| Sample ID | *MYBPC3*<br>on-target genotype | SNP14 genotype<br>(rs10769253, -2085 bp) | SNP15 genotype<br>(rs10769254, -1959 bp) |
| --- | --- | --- | --- |
| Egg donor 2 | WT/WT | G/G | G/G |
| MYBPC3 WT sperm donor | WT/WT | A/A | C/C |
| MYBPC3-M21.1 | homo-Indel/Indel | A/A | C/C |
| MYBPC3-M22.2 | homo-Indel/Indel | G/A | G/C |
| MYBPC3-M22.7 | homo-Indel/Indel | G/G | G/G |
| MYBPC3-M23.1 | homo-Indel/Indel | G/A | G/C |
| MYBPC3-M23.2 | homo-Indel/Indel | G/G | G/G |
| MYBPC3-M23.3 | homo-Indel/Indel | G/G | G/G |
| MYBPC3-M23.4 | homo-Indel/Indel | G/A | G/C |
| MYBPC3-M23.5 | homo-Indel/Indel | G/A | G/C |
| MYBPC3-M23.7 | homo-Indel/Indel | G/A | G/C |
| MYBPC3-M24.1 | homo-Indel/Indel | G/A | G/C |
| MYBPC3-M24.4 | homo-Indel/Indel | A/A | C/C |
| MYBPC3-M25.5 | homo-Indel/Indel | G/A | G/C |
| MYBPC3-M25.8 | homo-Indel/Indel | A/A | C/C |
| MYBPC3-M26.2 | homo-Indel/Indel | A/A | C/C |
| MYBPC3-M26.3 | homo-Indel/Indel | G/G | G/G |
| MYBPC3-M26.4 | homo-Indel/Indel | A/A | C/C |
| MYBPC3-M27.1 | homo-Indel/Indel | G/G | G/G |
| MYBPC3-M27.2 | homo-Indel/Indel | G/G | G/G |
| MYBPC3-M27.4 | homo-Indel/Indel | G/G | G/G |
| MYBPC3-M27.5 | homo-Indel/Indel | G/G | G/G |
| MYBPC3-M28.4 | homo-Indel/Indel | G/G | G/G |
| MYBPC3-M28.8 | homo-Indel/Indel | G/G | G/G |
| MYBPC3-M29.1 | homo-Indel/Indel | G/G | G/G |
| MYBPC3-M29.2 | homo-Indel/Indel | G/G | G/G |
| MYBPC3-M29.3 | homo-Indel/Indel | G/G | G/G |
| MYBPC3-M30.1 | homo-Indel/Indel | A/A | C/C |
| MYBPC3-M30.2 | homo-Indel/Indel | G/A | G/C |
| MYBPC3-M31.3 | homo-Indel/Indel | G/G | C/C |
| MYBPC3-M31.4 | homo-Indel/Indel | G/A | G/C |
| MYBPC3-M31.5 | homo-Indel/Indel | G/G | C/C |

